# The RNA-binding protein RBP10 controls a regulatory cascade that defines bloodstream-form trypanosome identity

**DOI:** 10.1101/076273

**Authors:** Elisha Mugo, Christine Clayton

**Affiliations:** DKFZ-ZMBH Alliance, Zentrum für Molekulare Biologie der Universität Heidelberg, University of Heidelberg. Im Neuenheimer Feld 282, D69120 Heidelberg, Germany.

## Abstract

Gene expression control in the pathogen *Trypanosoma brucei* relies almost exclusively on post-transcriptional mechanisms, so RNA binding proteins must assume the burden that is usually borne by transcription factors. *T. brucei* multiply in the blood of mammals as bloodstream forms, and in the midgut of Tsetse flies as procyclic forms. We show here that a single RNA-binding protein, RBP10, defines the bloodstream-form trypanosome differentiation state. Depletion of RBP10 from bloodstream-form trypanosomes gives cells that can grow only as procyclic forms; conversely, expression of RBP10 in procyclic forms converts them to bloodstream forms. RBP10 binds to procyclic-specific mRNAs containing an UAUUUUUU motif, targeting them for translation repression and destruction. Products of RBP10 target mRNAs include not only the major procyclic surface protein and enzymes of energy metabolism, but also protein kinases and stage-specific RNA-binding proteins: consequently, alterations in RBP10 trigger a regulatory cascade.

## Introduction

In multicellular organisms, cell differentiation is usually unidirectional, with stabilisation of differentiated states by epigenetic mechanisms, and reversal only in when healthy gene regulation is disrupted. In contrast, differentiation in unicellular organisms can be reversible, or involve cycles in which the direction of each transition is fixed. In each case differentiation is initiated by environmental stimuli, which activate signal transduction cascades; these in turn result in post-translational modifications that modify transcription factor activity. The consequent changes in mRNA synthesis, and hence protein synthesis, trigger a cascade of downstream regulatory events, affecting additional signalling molecules, transcription factors, and post-transcriptional modulators. Differentiation states are then stabilised by modification of DNA and chromatin.

Kinetoplastids are unicellular flagellates, and include many pathogens of mammals and plants, including the leishmanias and trypanosomes that cause human disease. Their life cycles involve a series of unidirectional transitions, which are driven by environmental changes as they move between mammalian and invertebrate hosts. The kinetoplastid studied in this paper, *Trypanosoma brucei*, causes human sleeping sickness in Africa and diseases of ruminants throughout the tropics. *T. brucei* multiply as bloodstream form trypomastigotes in the blood and tissue fluids of mammals, escaping immunity by antigenic variation of Variant Surface Glycoprotein (VSG) [1].

High cell density triggers growth arrest and differentiation to stumpy forms [2], which, upon uptake by a Tsetse fly, convert to proliferative procyclic forms in the midgut.

Procyclic forms express EP and GPEET procyclin surface proteins [3], and have numerous metabolic adaptations including increased mitochondrial metabolism. In the following 2 weeks, trypanosomes migrate towards the salivary glands [4].

Epimastigotes, expressing the surface protein BARP [5], are found attached to the proventriculus and salivary glands; here, a subset of parasites undergoes meiosis and mating [6,7]. Finally, growth-arrested, VSG-expressing metacyclic forms are seen swimming freely in the salivary glands; these are now ready for infection of a new mammal. Trypanosome developmental transitions are marked by many changes in mRNA [8-12] and protein [13-16] abundance.

Despite extensive changes in gene expression through the *T. brucei* life cycle, and the existence of stable differentiation states, the normal paradigm for regulation does not apply. This is because - as in other Kinetoplastid protists-transcription by *T. brucei* RNA polymerase II is polycistronic, with no regulation at the level of individual mRNAs. Monocistronic mRNAs are excised by 5′ *trans* splicing of a 39nt capped leader sequence, and by 3′ polyadenylation [17,18]. Regulation is thus exclusively at the levels of mRNA processing, degradation, and translation [10-12,19-22]. The sequences required for regulation of translation and mRNA decay often lie in the 3′-untranslated regions (3′-UTRs) of the mRNAs. Many groups of trypanosome mRNAs are coordinately regulated [23] and in a few cases, co-regulated genes have been shown to share regulatory elements that are bound by specific RNA-binding proteins [24,25]. RNA-binding proteins thus assume the burden that is carried by transcription factors in other eukaryotes, by regulating distinct mRNA subsets [19,26]. Although an RNAi machinery exists, it is not essential for cell proliferation or differentiation [27], and microRNAs have not been found [28].

The very abundant *VSG* and procyclin mRNAs are made, exceptionally, by RNA polymerase I [29], and their transcription is developmentally regulated by epigenetic mechanisms [1]. Procyclin is not subject to antigenic variation, and the presence of even a small amount on the bloodstream-form surface would enable the development of host immunity. Suppression of procyclin expression is therefore essential for trypanosome pathogenicity. However, procyclin gene transcription is only 10-fold less active in bloodstream forms than in procyclic forms [30]. This means that additional suppressive mechanisms are required: the mRNAs are both extremely rapidly degraded and very poorly translated [31-35], and trafficking of procyclin proteins to the surface is suppressed [36]. A 26nt motif within the 3′-UTR of procyclin mRNAs is required for their degradation and translational suppression [33,34]. However, until now, the protein(s) responsible for preventing procyclin expression were unknown.

RBP10 is a cytosolic protein with a single RNA recognition motif. Proteome analyses [16] and Western blot results [37] indicate that RBP10 is expressed in proliferating bloodstream forms, but not in stumpy forms or procyclics; *RBP10* mRNA also has low abundance in procyclic forms and in salivary gland trypanosomes [38]. Depletion of RBP10 by RNA interference (RNAi) in the bloodstream form is lethal, and forced RBP10 expression in procyclics inhibits growth [37]. Microarray analyses on cells with RBP10 RNAi or forced expression suggested a function for RBP10 either in stabilizing bloodstream-form specific mRNAs, or destabilizing procyclic-specific mRNAs, but it was unclear which effects were due to growth inhibition and the direct RNA targets of RBP10 were unknown.

In this paper we set out to identify the direct mRNA targets of RBP10 and to determine its role in mRNA control. We discovered that the mRNAs that are bound to RBP10 in bloodstream forms are preferentially targeted for translation inhibition and destruction, and that cells without RBP10 can survive only by differentiating to the procyclic form. Conversely, cells expressing RBP10 can survive only as bloodstream forms. The presence of RBP10 thus defines bloodstream-form trypanosome identity.

## Results

### Attachment of the RBP10 C-terminal domain to a reporter mRNA causes translation inhibition and mRNA degradation

Inducible expression of lambdaN-peptide-tagged RBP10 in cells containing a *CAT-boxB* reporter will result in attachment of the RBP10 to the reporter RNA via the lambdaN-boxB interaction. We previously showed that after 24h lambdaN-RBP10 induction, there was a decrease in boxB-tagged chloramphenicol acetyltransferase (*CAT*) reporter expression [37,39,40]. New deletion analyses (Fig 1) revealed that the C-terminal 88 amino acids of RBP10 was sufficient for the repressive effect (Fig 1 A, fragment F3). To investigate the mechanism in more detail we examined whether the *CAT-boxB* mRNA comigrated with polysomes in sucrose gradients. In the absence of tethered protein, the *CAT-boxB* reporter migrated in the dense sucrose fraction with large polysomes (Fig 1 B). With lambdaN-RBP10-F3 expression, *CAT-boxB* mRNA was reduced in abundance and the remainder migrated towards the top of the gradient. The control mRNA (encoding alpha-tubulin) was unaffected (Fig 1 B). The RBP10 C-terminus thus prevents translation initiation and promotes mRNA destruction. Interestingly, although a homologue of RBP10 is present in all sequenced Kinetoplastids, only the RRM domain is conserved: this C-terminal sequence is specific to *Trypanosoma* (S1 Fig).

**Fig 1.**
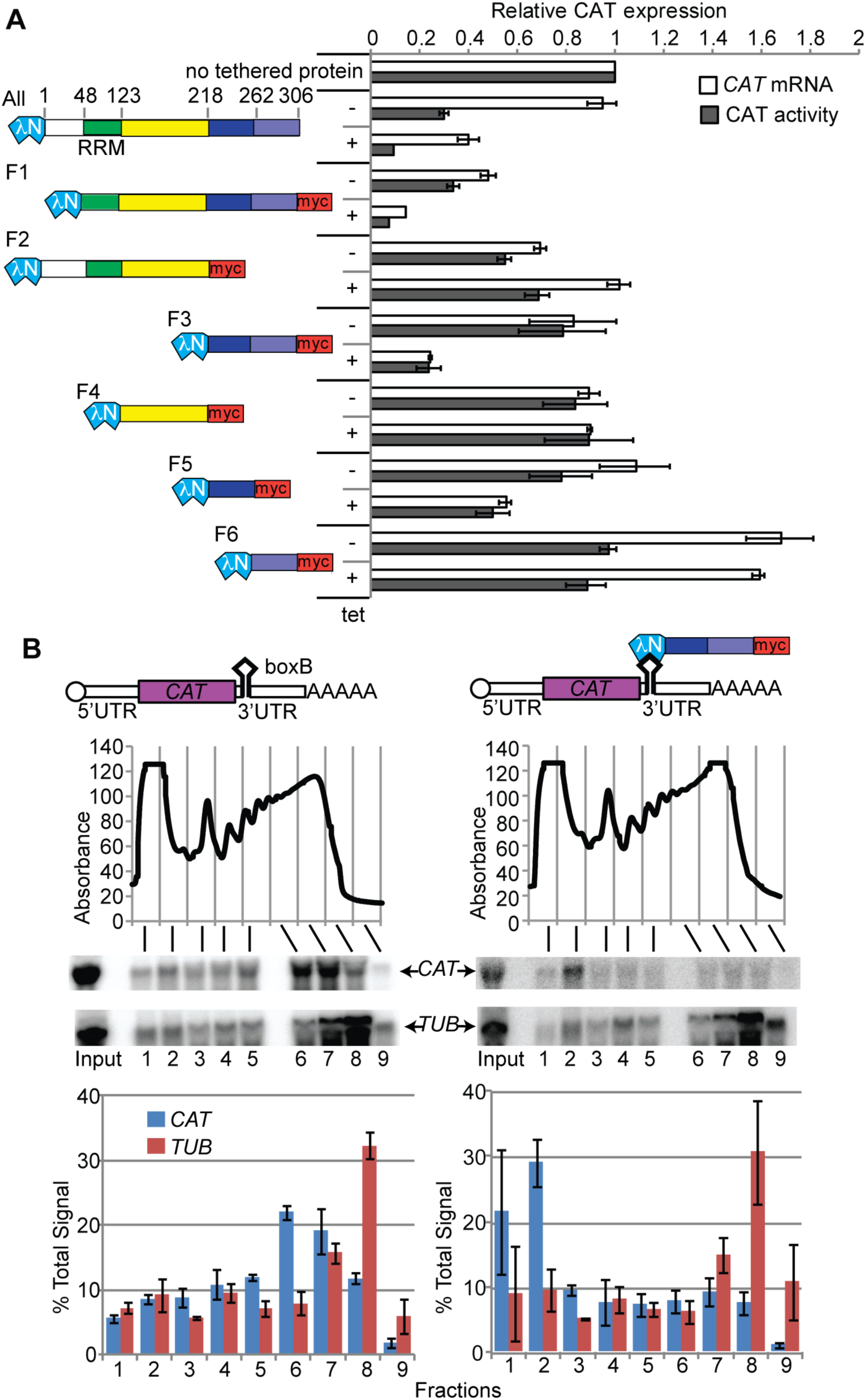
Tethering of the C-terminal portion of RBP10 to a reporter mRNA inhibits its translation and causes its destruction. A. Various fragments of RBP10 were inducibly expressed with an N-terminal lambdaN peptide and a C-terminal myc tag, in a cell line expressing a *CAT* reporter mRNA with 5 boxB sequences in the 3′-UTR. Cartoons showing the fragments are on the left. The numbers above the full-length protein are amino acid residues. The effects on the amounts *CAT* mRNA and CAT protein were measured by Northern blot and enzyme assay, respectively. Levels are expressed as mean ± standard deviation for at least 3 replicates, relative to a line with no lambdaN peptide protein (top bars) B. Bloodstream-form trypanosomes expressing the reporter in (A), and with tetracycline-inducible expression of lamdaN-RBP10-F3, were used. The reporter with or without tethered protein is shown at the top. Lysates from cells grown without (left) or with (right) tetracycline were subjected to sucrose gradient centrifugation. The panel below the cartoons show the absorption profiles at 254 nm as the fractions were collected. The migration of *CAT* and beta-tubulin (*TUB*) mRNAs on the gradient was detected by Northern blotting; representative blots are shown. The graphs show Northern signals, expressed as the percentage of the total signal, with arithmetic mean and standard deviation for three independent biological replicates.

### RBP10 can interact with an eIF4E interacting protein

We investigated the mechanism of RBP10-mediated repression by looking for interaction partners. We made a bloodstream-form cell line containing a single copy of RBP10 bearing a cleavable N-terminal affinity purification tag (S2 Fig A). After tandem affinity purification the most consistent co-purifying proteins were components of the proteasome (S1 Table). While this could be a mechanism for repression, it is also possible that some of the affinity-tagged RBP10 was targeted to the proteasome because it was incorrectly folded. The only interaction partner linked to mRNA degradation was CAF40, a component of the NOT deadenylase complex, but since no other components of the complex were present the significance of the interaction was unclear. Furthermore, V5-CAF40 pull-down failed to identify any RBP10 peptides (data not shown). To find out whether other proteins - including those of unknown function - might be important we investigated direct interactions of RBP10 by 2-hybrid screening. Nearly 80 potential partners were seen (S2 Table), of which 40 had not been seen in other screens. This suggested that RBP10 is prone to non-specific interactions. Potential partners included RBP10 itself, and four other RNA binding proteins. Interestingly, 8 of the 33 specifically interacting proteins had previously been show to suppress expression in the tethering assay [39,40]. There was however no overlap between the two-hybrid results and those from affinity purification.

A single fragment of a trypanosome eIF4E-interacting protein (4EIP) was scored as a possible RBP10 partner in the two-hybrid screen. Leishmania 4EIP interacts with eIF4E1 [41], which is one of the six trypanosomatid homologues of the cap-binding translation initiation factor eIF4E. In *T. brucei*, both 4EIP and eIF4E1 are extremely strong suppressors of reporter expression in the tethering assay [39,40]. We confirmed the two-hybrid interaction between full length RBP10 and 4EIP (Fig 2 A); and found that the interaction was mediated by the C-terminal 88 amino acids (F3) that are active in the tethering assay. This would be consistent with RBP10 acting via 4EIP recruitment. To test the interaction *in vivo*, we used trypanosomes expressing RBP10 that was N-terminally V5-tagged at the endogenous locus, then expressed C-terminally myc-tagged 4EIP from a strong promoter. Immunoprecipitation of myc-tagged 4EIP indeed pulled down a very small proportion of V5-RBP10 (Fig 2 B). It is therefore possible that RBP10 acts by recruiting 4EIP. However, depletion of 4EIP does not reproducibly inhibit bloodstream-form trypanosome growth ([42] and our preliminary results), which argues against this hypothesis.

**Fig 2.**
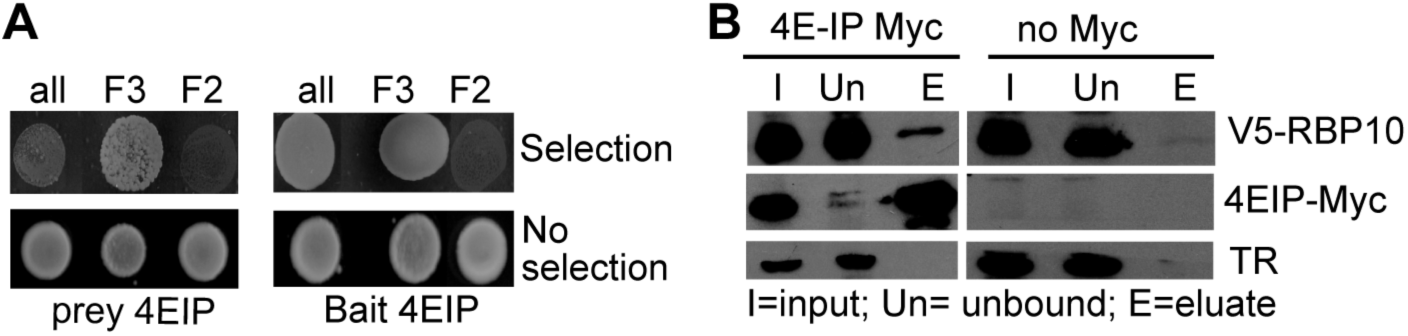
Interaction of RBP10 with 4E-IP A. Two-hybrid assay showing yeast growth. Upper panels show selection against non-interacting pairs. B. Immunoprecipitation of 4E-IP-Myc pulls down V5-tagged RBP10. The number of cell equivalents loaded for the eluate was 33x that of input and unbound fractions. The inverse precipitation did not confirm the interaction, but this could be because of an imbalance of protein abundances in the cells. (The abundance of 4E-IP is not known).

### RBP10 represses procyclic-form specific mRNAs

To analyse the role of RBP10 in gene expression in bloodstream forms, we depleted RBP10 by RNAi for 15h (S2 Fig B), before parasite proliferation and translation (S3 Fig) were affected. High-throughput cDNA sequencing (RNA-Seq) revealed nearly 200 mRNAs that changed significantly in total and/or polysomal RNA fractions (S3 Table, sheets 3-7). 36% of the mRNAs that increased after RNAi were preferentially expressed in procyclic forms, while 44% of those that decreased were preferentially expressed in bloodstream forms (Table 1 and Fig 3 A). Increases in polysomal RNA after RNAi were very slightly stronger than for total RNA. *RBP10* was the only mRNA with strongly decreased polysomal loading after RNAi, whereas about 60 mRNAs moved into polysomes (S3 Table sheet 3). 19 of these 60 mRNAs have a higher ribosome density in procyclic forms than in bloodstream forms [11], suggesting that their translation is inhibited in bloodstream forms. An example is the *COXVI* mRNA (S3 Table sheet 3, S4 Fig A,B), which is known to be developmentally regulated mainly at the level of translation [43]. Only 4 mRNAs had lower polysome occupancy after RNAi (S1 Table sheet 3). Our results almost certainly under-estimate the extent of translation changes, since we measured only whether an mRNA was in the polysomal fraction or not. For example, if an mRNA had a few ribosomes before RNAi, and more ribosomes after RNAi, this would not have been detected.

**Fig 3.**
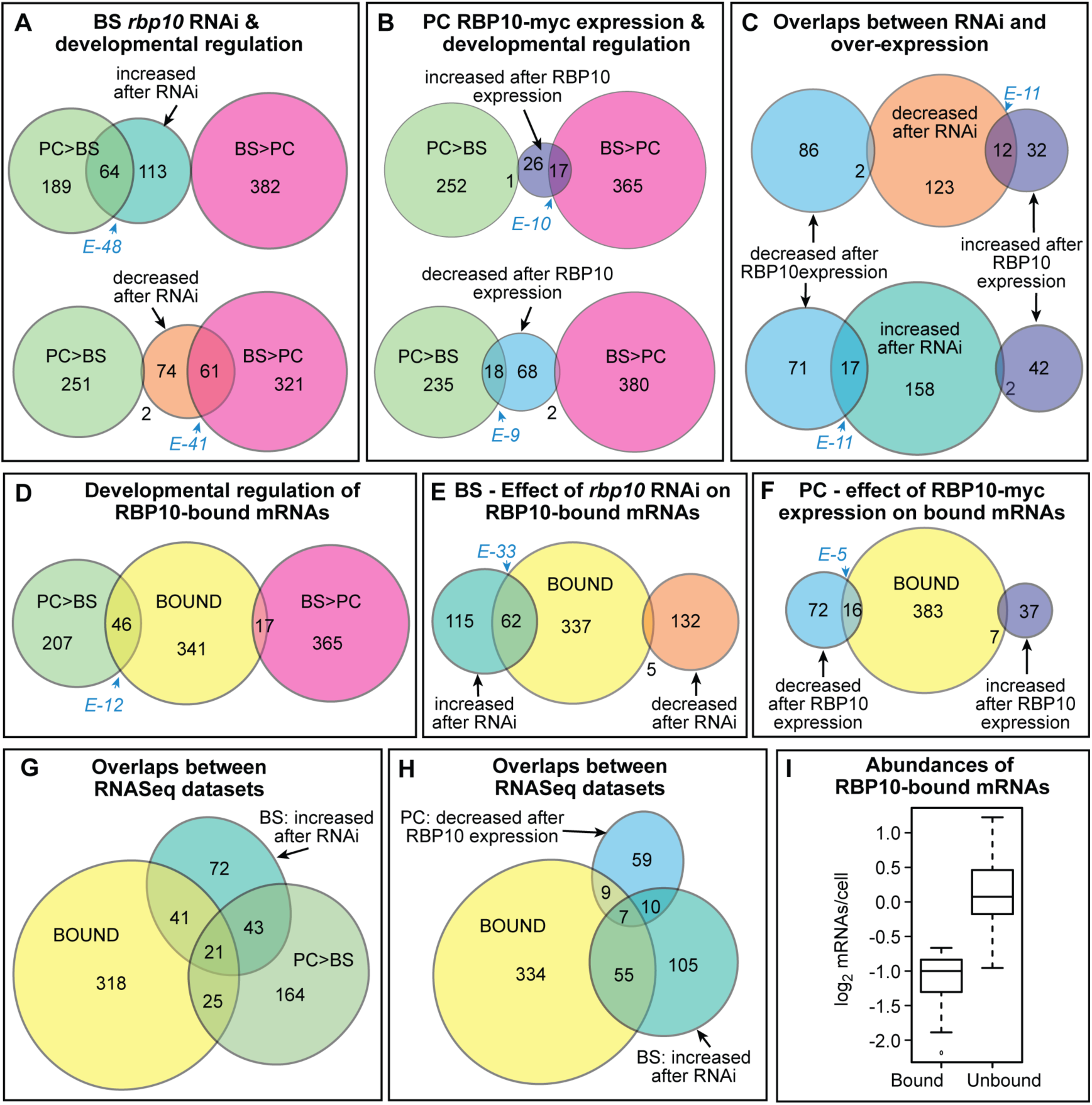
mRNAs bound to and/or regulated by RBP10. Results for mRNA levels are displayed as proportional Venn diagrams made using http://jura.wi.mit.edu/bioc/tools/venn.php and http://www.eulerdiagrams.org/eulerAPE/ BS = Bloodstream form, PC = procyclic form. Categories were: Bound: Supplementary Table S3 sheet 4. PC>BS and BS>PC: at least 2x significantly regulated [10]. BS rbp10 down or up: affected by RNAi, Supplementary Table S1 sheets 6 & 7; PC+RBP10 down or up: affected by RBP10 expression in procyclics, Supplementary Table S2 sheets 6 & 7. Fischer exact test values, calculated for the dataset in the centre, are in blue italics. A) Relationship between developmental regulation and the effects of RBP10 depletion in bloodstream forms B) Relationship between developmental regulation and the effects of RBP10 expression in procyclic forms C) Expression of RBP10 in procyclic forms related to the effects *rbp10* RNAi in bloodstream forms. D) Developmental regulation of the mRNAs bound by RBP10. E) Effect of bloodstream-form *rbp10* RNAi on the mRNAs bound by RBP10. F) Effect of procyclic-form RBP10 expression on the mRNAs bound by RBP10. G) Overlaps between bound mRNAs, procyclic-specific mRNAs, and the mRNAs that increased after *rbp10* RNAi in bloodstream forms H) Overlaps between bound mRNAs, the mRNAs that increased after *rbp10* RNAi in bloodstream forms, and the mRNAs that decreased after RBP10 expression in procyclic forms I) Abundances of RBP10-bound mRNAs compared to all other mRNAs.

Next we analysed the effect of expressing C-terminally myc-tagged RBP10 (RBP10-myc) in procyclic forms. After only 6h induction, (S2 Fig C, S4 Fig) we saw decreases in procyclic-form-specific mRNAs, and increases in bloodstream-form specific mRNAs (Fig 3 B and S4 Table). A subset of mRNAs showed opposite reactions in the bloodstream RNAi and procyclic over-expression experiments, consistent with being immediate RBP10 targets (Fig 3 C).

We had previously attempted to identify RBP10-bound mRNAs, but the methods that we used had insufficient sensitivity [37]. Since a subsequent study showed that RBP10 is indeed bound to mRNA [40], we made another attempt. This time, we used our bloodstream-form cell line in which all RBP10 bears a cleavable tag (S2 Fig A), and compared bound and unbound mRNAs by RNASeq. 400 mRNAs were identified (S5 Fig A and S5 Table, sheet 3). 12% of them are more abundant in procyclic forms than in bloodstream forms (Fig 3 D). Consistent with the tethering results, 15% of the RBP10 targets increased after *rbp10* RNAi in bloodstream forms, but only 1% decreased (Fig 3 E). Only 16 target mRNAs decreased after RBP10 expression in procyclic forms (Fig 3 F); this low number might be due to the very short duration of RBP10-myc expression. The RBP10-bound mRNAs that are more abundant in procyclic forms and increased after *rbp10* RNAi (Fig 3 G) are listed in Table 1: six of them are also decreased after RBP10 expression in procyclics (Fig 3 H). Scatter plots showing the effects of RNAi on bound RNA abundance and polysomal loading are shown in S5 Fig C and D.

**Table 1.** 21 mRNAs that are bound to RBP10, are more abundant (RNA PC>BS) or better translated in procyclic forms than in bloodstream forms according to ribosome profiling (Riboprof PC>BS), and are increased after *rbp10* RNAi in bloodstream forms. For details see S5 Table sheet 8. The latter includes many additional mRNAs that fulfilled fewer conditions.

| id | Annotation | RNA<br>PC><br>BS | RiboP<br>rof<br>PC>B<br>S | de-<br>creased<br>by RBP10<br>in PC |
| --- | --- | --- | --- | --- |
| **Metabolism** |  |  |  |  |
| 9.5900 | glutamate dehydrogenase (GDH) | yes | yes | no |
| 10.2350 | pyruvate dehydrogenase complex | yes | yes | no |
| 10.4280 | Complex III subunit | yes | yes | no |
| 1.4100 | cytochrome oxidase subunit IV (COXIV) | yes | yes | no |
| 11.15550 | NADH-cytochrome b5 reductase, | yes | yes | no |
| 4.4990 | ubiquinol-cytochrome c reductase | yes | yes | no |
| 7.210 | proline dehydrogenase | yes | yes | no |
| 9.4310 | tricarboxylate carrier | yes | yes | no |
| 9.7470 | purine nucleoside transporter NT10 | yes | yes | yes |
| 11.6280 | pyruvate phosphate dikinase (PPDK) | yes | yes | no |
| 11.8990 | cation transporter, putative | yes | yes | no |
| Surface |  |  |  |  |
| 10.10250 | Procyclins # | yes | yes | yes |
| 7.6850 | trans-sialidase (TbTS) | yes | yes | yes |
| 8.7340 | trans-sialidase | no | yes | yes |
| 4.3500 | Amastin-like protein | yes | yes | no |
| **RNA binding** |  |  |  |  |
| 7.2660 | ZC3H20 | yes | no | no |
| 7.2680 | ZC3H22 | yes | yes | yes |
| **Unknown function** |  |  |  |  |
| 9.13200 | hypothetical protein | yes | yes | yes |
| 11.16300 | hypothetical protein | yes | yes | no |
| 7.5550 | hypothetical protein | yes | yes | no |
| 10.3300 | DUF4206 (cysteine-rich) | yes | yes | no |
| 10.2360 | hypothetical protein | yes | no | no |

Overall, the RBP10-bound mRNAs are significantly less abundant than unbound mRNAs (Fig 3 I). At the proteome level [16], the products of RBP10-bound mRNAs are enriched in proteins that increase within 2-24h of the initiation of stumpy form differentiation, or early during conversion to procyclic forms. There is no enrichment for proteins that appear later in differentiation, or only in mature procyclics, and unregulated proteins or those that are decreased in procyclic forms are under-represented (S5 Table, sheet 6).

### RBP10-bound mRNAs share a UAUUUUUU motif

A comparison of the 3′-untranslated region (3′-UTR) sequences of the most reliably RBP10-bound mRNAs with those of unbound mRNAs (S5 Table, sheet 5 and S1-S3 texts) revealed that the motif UA(U)_6_ is highly enriched in the 3′-UTRs of RBP10 targets (Fig 4 A). It is present 1-9 times in 225 out of the 255 most strongly bound 3′-UTRs (Fig 4 B, S1 text). Transcriptome-wide, the number of UA(U)_6_ motifs was strongly associated with RBP10 binding (Fig 4 C). Nevertheless, many mRNAs that contain UA(U)_6_ were not pulled down with RBP10. Although there are some exceptions [44,45], RRM domains usually bind to single-stranded RNA [46-48]; the unbound motifs may have inappropriate conformations, or they might instead be bound by competing proteins.

**Fig 4.**
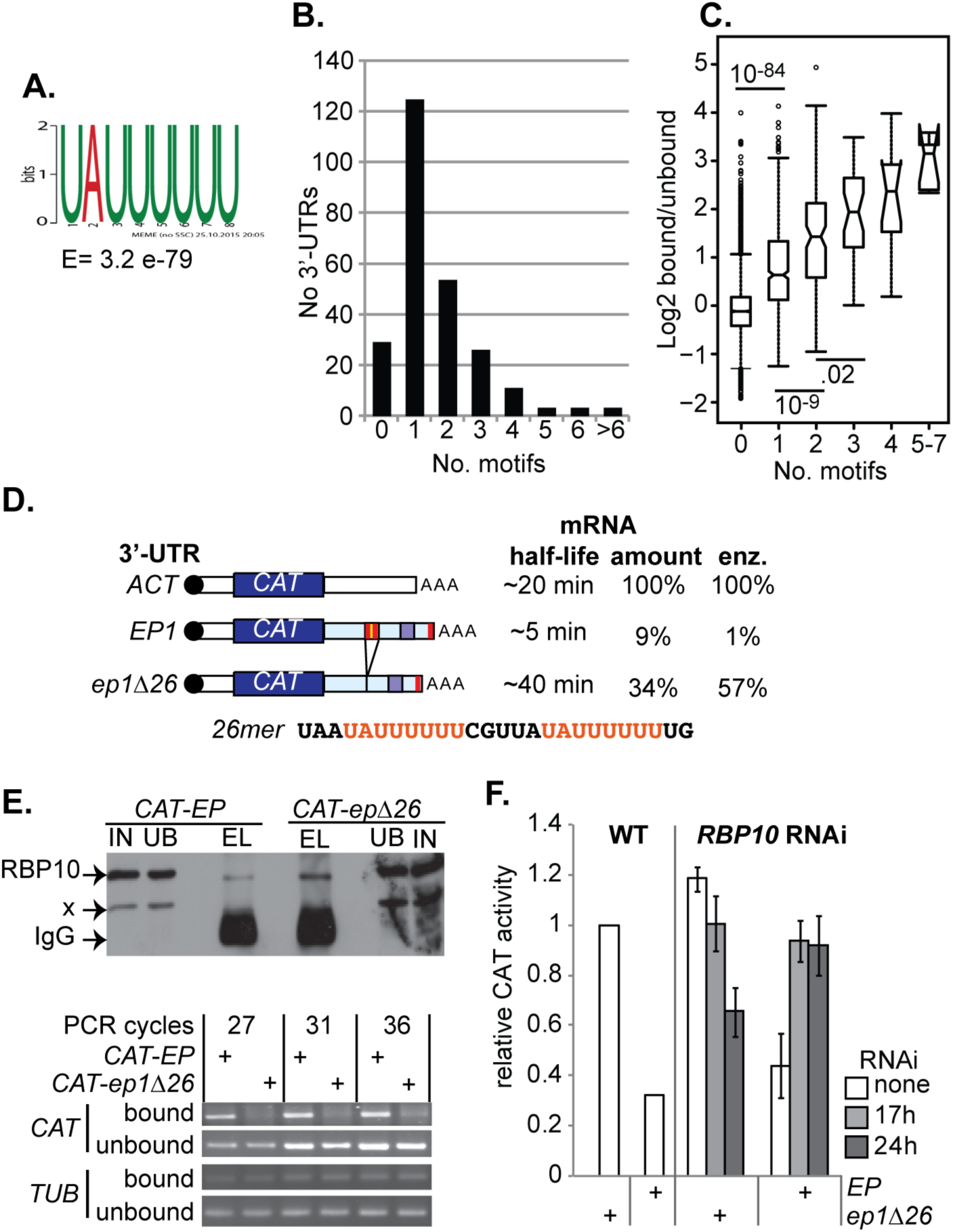
RBP10 targets a UAUUUUUU motif A) The 3′-UTRs of 188 mRNAs that were at least 3x enriched in both RBP10-bound samples (Supplementary Table S5 sheet 5) were compared with those of mRNAs that did not bind (<0.7x enrichment in both experiments) using DREME. Only 3′-UTRs annotated in tritrypDB were used (S1 text, S2 text, S3 text). The best-scoring motif found is shown. B) Numbers of UA(U)_6_ motifs in 255 manually annotated bound 3′-UTRs (S1 text, S5 Table sheet 5). One mRNA each had 7, 8 and 9 motifs. The numbers of motifs shown here are sometimes higher than with automatically annotated 3′-UTRs from the genome database, because some of the automatic 3′-UTRs are truncated. C) Effect of the UA(U)_6_ motif on RBP10 binding: median ± 25th percentiles, with 95% confidence limits and outliers. Database 3′-UTRs were analyzed. Since these are often truncated, and are missing for 30% of genes, the numbers of motifs are under-estimated. Significant differences (Student t-test) are shown, For the mRNAs that were enriched according to DeSeq (S5 Table, sheet 3), 284/353 annotated UTRs contained UA(U)_6_. D) Reporters used in (E) and (F) [34]. The red bars are UAUUUUUU elements, which are highlighted in the 26mer sequence. E) RBP10 binding to the *EP* 3′-UTR requires the 26mer. Cells were UV-irradiated, and RBP10 was pulled down with anti-RBP10 (upper panel, Western blot). RNAs were detected by reverse transcription and PCR using gene-specific primers (lower panel). F) *rbp10* RNAi specifically affects expression of a reporter containing the 26mer. CAT activity was measured 24h after RNAi induction.

UA(U)_6_ was previously identified in several mRNAs encoding procyclic-specific proteins [43]. The 26mer 3′-UTR element that is required for *EP1* procyclin mRNA instability [33,34] contains two UA(U)_6_ motifs (Fig 4 B) and these are single-stranded *in vivo* [49], as is required for RRM domain binding. Deletion of the region containing UA(U)_6_ from the COXV 3′-UTR increased protein expression without much effect on the mRNA level [43], consistent with translation control. The mRNAs encoding phosphoglycerate kinase (PGK) constitute a third example. In bloodstream-form trypanosomes, most PGK is encoded by the gene *PGKC* and is targeted to microbodies called glycosomes (S6 Fig A); in procyclic forms, a different gene, *PGKB*, is expressed and the enzyme is in the cytosol [50,51] (S6 Fig C). The *PGKB and PGKC* coding regions are identical except for a C-terminal targeting sequence in *PGKC*, so they were not distinguished in our initial RNASeq analysis. However, the 3′-UTRs are completely different, and determine developmental regulation [52]. The UA(U)_6_ motif is present twice within a 3-UTR segment that is required for suppression of PGKB expression in bloodstream forms [53] (S1 text). Using our RNASeq pull-down data, we compared the abundances of several 15nt sequences specific to the *PGKB* and *PGKC* mRNAs and found that that *PGKB* mRNA is almost exclusively in the bound fraction, whereas most *PGKC* mRNA is unbound. A similar analysis gave RBP10 binding ratios of 2.8:1 for *EP1* and 1.3:1 for *GPEET* mRNAs. The *GPEET* mRNA 3′-UTR has UA(U)_5_ repeats, but no UA(U)_6_, and it is not yet clear whether it is bound by RBP10.

To test whether the 26mer was required for binding of the *EP1* 3′-UTR to RBP10, we used cell lines that expressed chloramphenicol acetyltransferase (CAT) reporter mRNAs with either the intact wild-type *EP1* 3′-UTR, or a version without the 26mer (*ep1∆26*) (Fig 4 D) [34]. Co-immunoprecipitation of the *CAT* mRNA with RBP10 depended on the presence of the 26mer (Fig 4 E). Depletion of RBP10 by RNAi also increased expression from the *CAT-EP* reporter but not from *CAT-ep1∆26* (Fig 4 F). These results demonstrated both that RBP10 binds to the *EP* 3′-UTR via the 26mer sequence, and that the 26mer is necessary for RBP10-mediated regulation of an mRNA containing the *EP1* 3′-UTR.

### Expression of RBP10 determines bloodstream form identity

To test the role of RBP10 in differentiation, we first examined conversion from the bloodstream form to the procyclic form. Our monomorphic bloodstream forms with *rbp10* RNAi, or treated with the differentiation-inducer *cis* aconitate, are unable to multiply under procyclic culture conditions. We therefore used differentiation-competent trypanosomes (EATRO 1125, Antat1.1 strain). As expected [3], growth of these cells at high density in methyl cellulose for three days resulted in expression of the a-form marker PAD1 [54] and stumpy morphology; RBP10 was also reduced (Fig 5 A, lanes 2 &3), consistent with its absence in the stumpy-form proteome [54]. The cells were now treated with 6mM cis-aconitate in bloodstream-form medium at 27°C for 17h. Upon transfer to procyclic medium without cis-aconitate, the cells started to grow as procyclics within 24h (Fig 5B).

**Fig 5.**
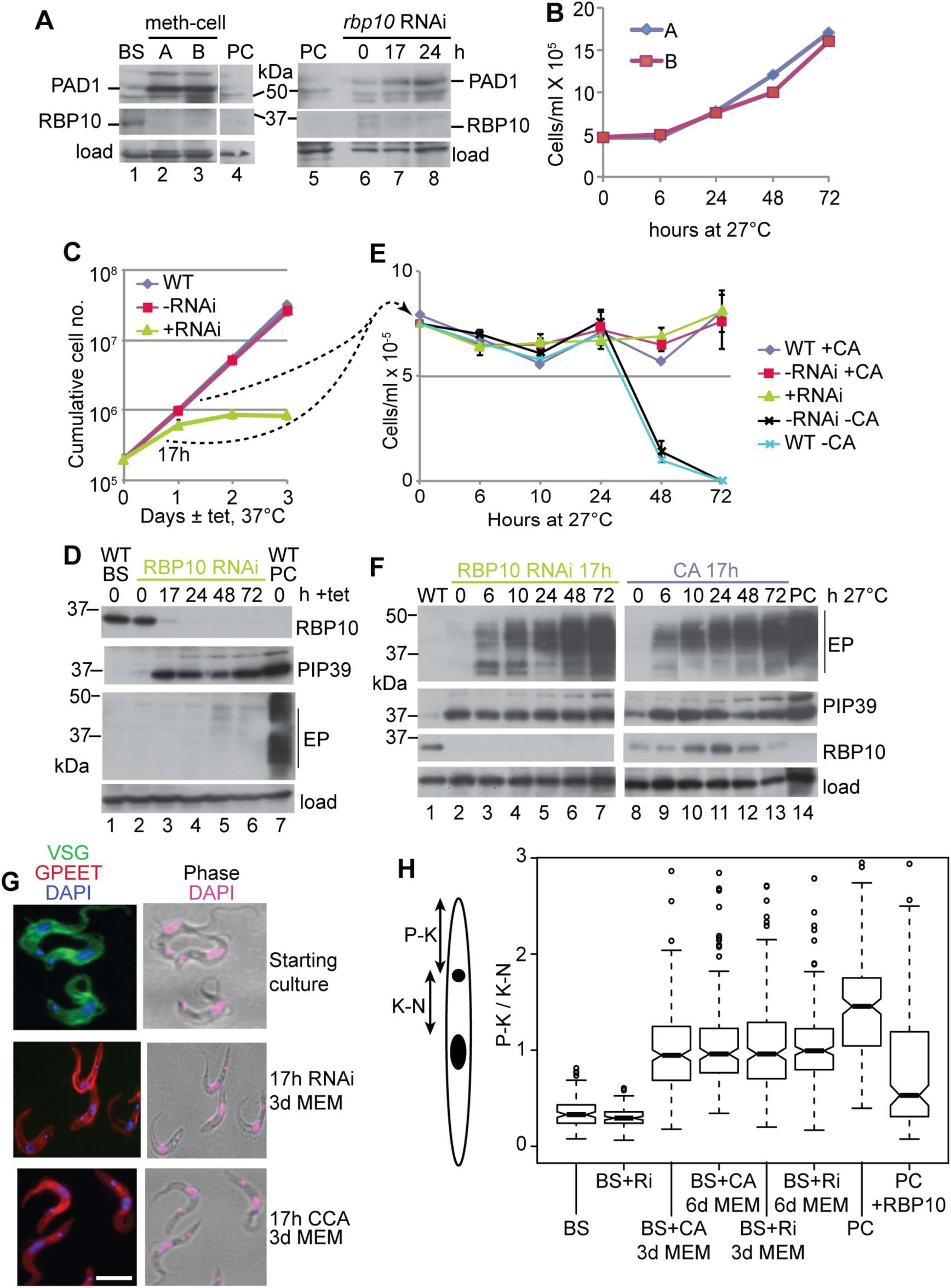
RBP10 depletion primes bloodstream forms for differentiation to procyclic forms A) Expression of PAD1 and RBP10 in trypanosomes incubated at maximal density in methyl cellulose-containing medium for 3 days (left panel), or after *rbp10* RNAi (right panel). The RBP10 signal is weak, probably because the samples were not denatured in order to allow PAD1 detection. B) Growth of trypanosomes incubated at maximal density in methyl cellulose-containing medium for 3 days, followed by cis-aconitate treatment for 17h at 27°C. Time 0 is the time of transfer to procyclic conditions. C) Cumulative growth curve of bloodstream-form (BS) trypanosomes with and without RNAi. WT = wild-type (*tet* repressor only);-RNAi: RNAi cell line with no tetracyline: +RNAi: RNAi cell line with tetracycline. D) Expression of RBP10, PIP39, and EP procyclin were measured by Western blotting up to 3d after tetracycline induction of RNAi. E) 17h after RNAi induction at 37°C in bloodstream form medium (+RNAi), or incubation at high density in procylic medium with 6mM cis-aconitate at 27°C (+CA), trypanosomes were placed in procyclic medium (MEM-pros) at 27°C with neither tetracycline nor cis-aconitate. The cell densities of the cultures were monitored. Negative controls were either wild-type or uninduced RNAi lines, without cis-aconitate. Cell densities are shown; they started to increase at 72h. F) Protein expression after transfer to procyclic medium at 27°C after 17h pre-treatment as in (E). G) Immunofluorescence: Cell morphology and expression of GPEET procyclin after 3d cultivation in procyclic medium. DNA is stained with DAPI. H) The cartoon shows the way in which the position of the kinetoplast, relative to the nucleus (P-K/K-N), was measured. The box plot shows the median with 25th percentiles and 95% confidence limits. Some outliers have been deleted. The full results, together with the numbers of trypanosomes measured, are in S7 Fig A.

BS - normal bloodstream forms; PC- normal procyclic forms; CA - 17h cis-aconitate pretreatment; RNAi or Ri; 17h *rbp10* RNAi induction; +RBP10 - induced expression of RBP10 for 2 days. There were no significant differences between the 3-day and 6-day differentiating populations. All other pairs were significantly different (Student t-test, p<0.05).

Induction of *rbp10* RNAi in differentiation-competent cells in log phase resulted in growth inhibition (Fig 5 C), weak PAD1 expression (Fig 5 A, lanes 6-8) and, within 17h, expression of the differentiation-regulating phosphatase PIP39 [55] (Fig 5 D). After 17h RNAi, transfer of the cells to procyclic-form medium at 27°C (without cis-aconitate) allowed them to survive and eventually start to grow (Fig 5 E), with expression of EP procyclin (Fig 5 F, lanes 1-7) and GPEET procyclin (Fig 5 G) proteins. As expected, cells without RNAi induction died (Fig 5 E). The RNAi experiment had to be done without high density pre-incubation in methyl cellulose since RNAi can only be induced in growing trypanosomes. As a control, we therefore grew differentiation-competent cells to maximum density in liquid medium, and treated them with cis-aconitate for 17h at 27°C [56] before transfer to procyclic medium without cis-aconitate, also at 27°C. In these cells, weak RBP10 expression persisted for 2 days (Fig 5 F, lanes 8-13) but the growth and procyclin expression kinetics were similar to those seen after *rbp10* RNAi (Fig 5 E, F, G).

The rather slow resumption of growth after both *rbp10* RNAi and 17h cis-aconitate treatment, and the persistence of RBP10 in the cis-aconitate experiment, suggested that the population might be a mixture of differentiation-competent and incompetent cells. Indeed, both differentiating cultures were mixtures of normal-looking, proliferating, and dying or abnormal forms, and even after 3 days in procyclic culture, about 9% of the cells in both cultures retained bloodstream-form morphology and PGK localization (S6 Fig D). In bloodstream forms, the kinetoplast is at the posterior end of the cell (Fig 5 G), which means that the distance between the posterior and the kinetoplast (P-K) is much less than the distance from the kinetoplast to the nucleus (K-N) (Fig 5 H and S7 Fig A, B; “BS”). Procyclic forms are longer and the kinetoplast is nearer to the nucleus (Fig 5 G), giving a much higher P-K/K-N ratio (Fig 5 H and S7 Fig A, B; “PC”). The differentiating cultures showed very heterogeneous P-K/K-N ratios, with medians that were intermediate between those of bloodstream and procyclic forms (Fig 5 H, S7 Fig A-C). As expected, cells with low ratios typical of bloodstream forms were preferentially GPEET negative (S7 Fig D,E).

We next inducibly expressed RBP10-myc in differentiation-competent procyclic forms. For the results shown, we made bloodstream forms with inducible RBP10-myc, differentiated them into procyclic forms, grew these for more than 3 months, and then induced RBP10 expression. However similar results were obtained by differentiating bloodstream forms into procyclic forms, and then making three new cell lines with the inducible RBP10-myc plasmid. After RBP10-myc induction, cell growth was inhibited, as expected [37] (Fig 6 A). After 48h, expression of procyclin was strongly reduced, alternative oxidase increased, and untagged RBP10 was detected (Fig 6 B, C). At this point the cell population was very heterogeneous. Some cells had normal procyclic morphology and expressed EP and GPEET procyclin; some were clearly abnormal; while another subset had clear bloodstream-form morphology (Fig 6D). This was reflected in the P-K/K-N ratios, which ranged from procyclic to bloodstream-form patterns (Fig 5 G, S7 Fig F). Flow cytometry analysis revealed that about 50% of cells had decreased, or absent, procyclin expression (Fig 6 E, F). Cells with bloodstream-form morphology were usually (but not always) procyclin negative (S7 Fig G). Notably, *VSG* mRNA was present (Fig 6 C). We do not know which VSGs are expressed in the bloodstream-form-morphology cells, but a subset of them stained with a polyclonal antibody against VSG117 (Fig 6 D, G, H). PGK staining in the bloodstream-form-like cells seemed reduced and somewhat more punctate than in procyclic forms, but had not yet assumed a purely glycosomal pattern (S6 Fig E). Remarkably, the RBP10-induced cells could now survive only if they were transferred to bloodstream-form growth conditions (Figure 6A). After 6 days in these conditions, all surviving cells had bloodstream-form morphology and 60% of them cross-reacted with the anti-VSG117 antibody.

**Fig 6.**
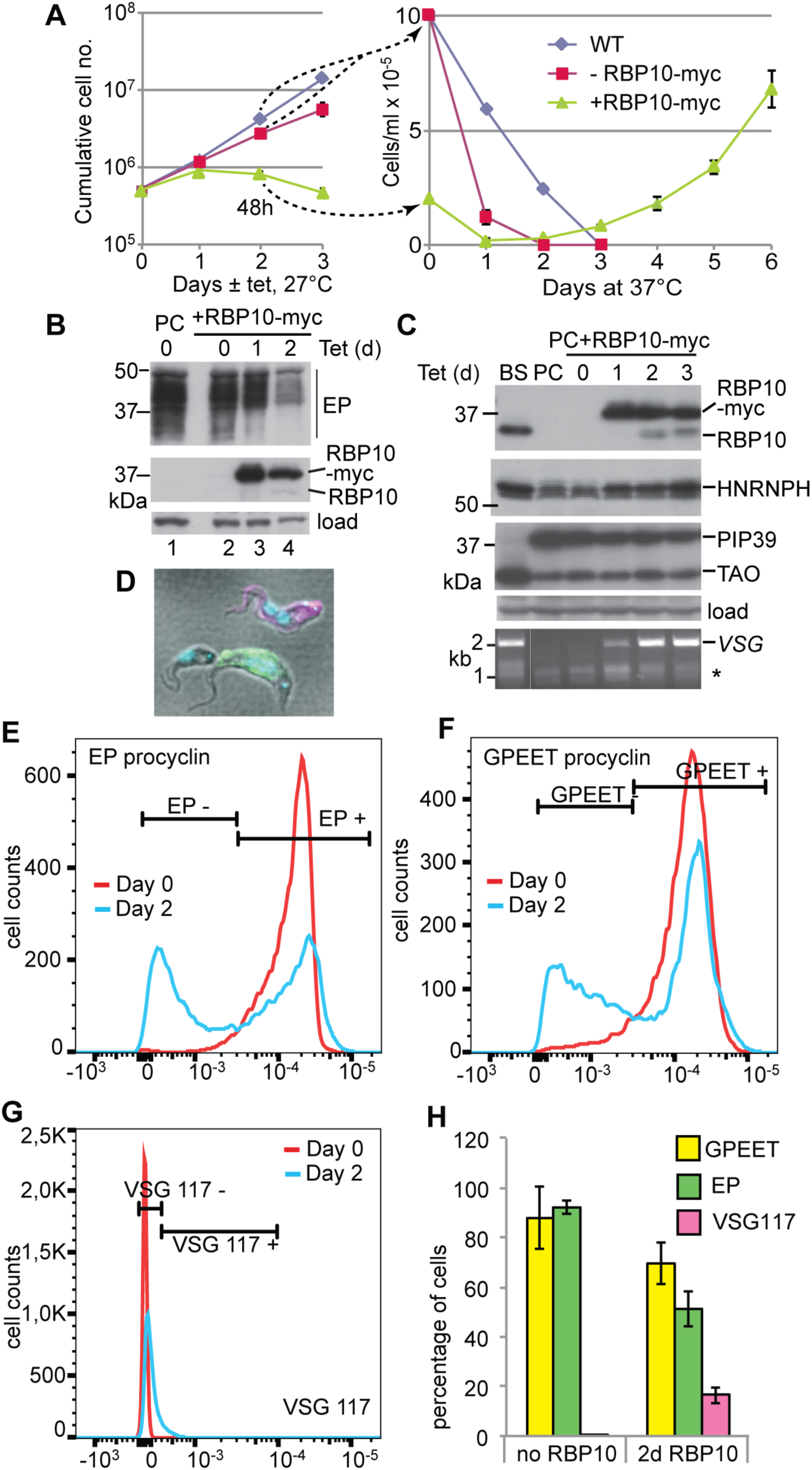
RBP10 expression in procyclic forms causes differentiation to bloodstream forms A) Cell counts in a typical experiment. Expression of RBP10-myc was induced using tetracycline at 27°C in procyclic-form medium. The left-hand panel is a cumulative growth curve. After 48h cells were transferred to bloodstream-form medium at 37°C; in the right-hand panel cell densities are shown. B) Expression of EP procyclin and RBP10 after induction of RBP10-myc expression. Cells were cultured in procyclic medium at 27°C. Expression of EP Procyclin protein decreased after 48h of RBP10-myc induction, while untagged RBP10 becomes detectable. C) As in (B) except that culture continued for 4 days. Proteins that are expressed more in bloodstream forms than in procyclic forms were detected by Western blotting. TAO is trypanosome alternative oxidase; HNRNPH is an RNA-binding protein. *VSG* mRNA was detected by RT-PCR using a spliced leader primer and a primer that hybridizes to a conserved region within the *VSG* 3′-UTR. D) Trypanosomes were taken after 48h RBP10-myc induction. They were stained for EP procyclin and phosphorylated GPEET procyclin (green), VSG117 (magenta) and DNA (cyan). The stain is overlaid with differential interference contrast (grey). The panel shows a typical green procyclic form, a bloodstream form with two kinetoplasts, and a third cell which resembles a bloodstream form (terminal kinetoplast) but was not stained by anti-VSG117. This cell may express a VSG that does not react with the anti-VSG117 antibodies. E) Quantitation of surface EP procyclin by flow cytometry, comparing cells with (day 2) and without (day 0) induced RBP10-myc expression. F) Quantitation of surface phospho-GPEET procyclin by flow cytometry, comparing cells with (day 2) and without (day 0) induced RBP10-myc expression. G) Quantitation of staining with anti-VSG117 by flow cytometry, comparing cells with (day 2) and without (day 0) induced RBP10-myc expression. H) Measurement of surface protein expression by FACSscan. Values for the different windows are mean and standard deviation from 4 replicates. Similar results were obtained by manual counting of stained smears.

The conversion of procyclic forms to bloodstream forms was highly reproducible. It was not due to persistence of a few bloodstream forms in the procyclic cultures, since survival always depended on induced RBP10 expression (Fig 6 A). RBP10 induction for two days was required; this was also the time needed to detect native RBP10 expression by Western blotting. Moreover, our preliminary results indicate that transient expression of RBP10 from an episome is sufficient to generate bloodstream forms. In contrast, attempts to obtain bloodstream forms by RBP10 expression in Lister 427 cells that have been in procyclic form culture for more than two decades were not successful.

Interestingly, epimastigote-like cells were never seen after induction of RBP10-myc expression in our differentiation-competent cells (24h or 48h), and the epimastigote-specific surface protein BARP was not detected by Western blotting or immunofluorescence. Although we may have missed transient formation of epimastigotes, this is unlikely since the behaviour of the trypanosomes was clearly not synchronous. Moreover, the presence of dividing cells with bloodstream-form morphology, suggested that RBP10 induction might be causing a direct conversion of procyclic forms to bloodstream forms, “jumping” past the non-dividing metacyclic form. We therefore compared the mRNA changes after 6h RBP10-myc induction with those found in salivary gland parasites (a mixture of epimastigotes and metacyclics) (S4 Table, sheets 5 and 6). The transcriptome comparisons and morphological analysis suggested that after 2 days of RBP10 expression the cell population might include both metacyclic and bloodstream forms. Interestingly, however, the mRNA encoding the meiosis protein MND1, which is specific to a pre-metacyclic intermediate form, is bound by RBP10 and *decreases* upon RBP10 induction. Evidence so far thus suggests that RBP10 induction causes a developmental switch that by-passes the epimastigote stage.

## Discussion

Our results indicate that the presence of RBP10 defines the identity of a trypanosome as a replication-competent bloodstream form.

RBP10 binds to procyclic-specific mRNAs with a UA(U)_6_ motif in the 3′-UTR, suppressing translation and causing mRNA destruction. However, many RNAs that are bound by RBP10 did not change in either abundance, or in the percentage in polysomes, after RBP10 RNAi or forced expression. As noted above, our assay did not measure ribosome density, so it is likely that many changes in translation went undetected. However, our results are completely consistent with work on other RNA-binding proteins: usually, only some of the bound mRNAs change after the protein is removed (e.g. [57]). For most mRNAs, many different proteins can bind along the same 3′-UTR - a sequence of 300nt is likely to bind at least twenty proteins - and it is their combinatorial effects that determine mRNA behaviour.

RBP10 target mRNAs include several that have been studied in considerable detail, including those encoding cytochrome complexes and the procyclins. However, the direct effects of RBP10 are by no means sufficient to explain its far-reaching influence, since many mRNAs that change in translation or abundance after RBP10 depletion are not bound by RBP10. This suggests that RBP10 depletion may initiate a cascade of events. Our results show how this could occur. Notably, the direct RBP10 targets include the mRNAs encoding three potential RNA-binding proteins, ZC3H20, ZC3H21, and ZC3H22, These proteins are normally expressed only in procyclic forms [15,16,58]. ZC3H20 binds to, and stabilises, at least two procyclic-specific mRNAs [58], of which one, *MCP12*, is not bound by RBP10 but was increased after *rbp10* RNAi. ZC3H22 suppresses expression in the tethering assay [39], and is required for proliferation of procyclic forms [15,42]. Suppression of ZC3H21 and ZC3H22 expression is clearly very important to bloodstream-form trypanosome survival: their mRNA 3-UTRs contain, respectively, no fewer than 5 and 7 RBP10 binding motifs.

In addition to controlling ZC3H20-22, RBP10 binds to and represses mRNAs encoding two procyclic-specific protein kinases (Tb927.8.6490, Tb927.11.15010) and one procyclic specific protein phosphatase (Tb927.10.8050). Also bound by RBP10, but with no detected effect of RNAi, are the transcripts encoding protein kinases RDK1 and NRKA. NRKA [15] is up-regulated after the onset of stumpy form differentiation; it is possible that RBP10 indeed regulates its translation but we did not detect this because of the insensitivity of our assay. RDK1 is a differentiation repressor [59] and is more abundant in bloodstream forms, making regulation by RBP10 unlikely. Similar to RBP10, depletion of *RDK1* primes bloodstream forms to differentiate to procyclic forms. More than 20 mRNAs show similar changes after RNAi targeting either *RDK1* or *RBP10*, and 16 of these are both bound and regulated by RBP10. Surprisingly, the *RBP10* transcript was not significantly affected after *RDK1* RNAi; however, the protein level and phosphorylation status of RBP10 were not determined.

The changes in RNA regulators, protein kinases and phosphatases, and perhaps also the changes in metabolism that are directly induced by RBP10 loss (e.g. [60]) will trigger downstream effects on signaling pathways. Such secondary effects of *rbp10* RNAi must include the increase in PIP39 (Tb927.9.6090), a key regulator of differentiation [55], and decreases in mRNAs encoding the RNA-binding proteins PUF11, RBP9, ZC3H31, ZC3H46, ZC3H48, DRBD5 and HNRNPH (S1 Table, sheet 7). These in turn will trigger further changes in gene expression.

So far, the only condition known to allow differentiation of *T. brucei* procyclic forms to bloodstream forms *in vitro* is induced expression of the putative RNA-binding protein RBP6. This mimics the natural differentiation pathway, with formation of epimastigotes within 24h and appearance of VSG-expressing metacyclic forms 4-5 days later [61]. *RBP6* mRNA has low abundance in both procyclic and bloodstream forms, and is maximal in salivary gland parasites [38]. RBP6 action is therefore transient, although it might possibly define the epimastigote form; its mechanism of action is unknown. Depletion of the RNA-binding protein DRBD18 from procyclic forms caused a partial switch to a bloodstream-form or epimastigote expression pattern; the mRNAs that increased included those encoding RBP6 and RBP10, but the ability to differentiate further was not tested [62]. Heat shock can also cause small increases in epimastigote-specific transcripts [63]. The rapid conversion of procyclic forms to bloodstream forms by RBP10 expression, in contrast, might be regarded as a form of *trans*-differentiation since the intermediate epimastigote stage was not detected. In animal cells, *trans*-differentiation can be achieved through changes in expression of transcription factors, either by direct manipulation or by changing expression of microRNAs that control transcription factor expression [64-66]. However, these changes have so far always been unidirectional. *Trans* differentiation that works in both directions according to the presence or absence of a single protein has, to our knowledge, not previously been observed.

It has long been known that stumpy forms cannot revert to long-slender bloodstream forms, and recent results indicate that the differentiation of stumpy forms to procyclics is an irreversible bi-stable switch [15]. We suggest that expression of RBP10 governs a bi-stable switch during the transition from bloodstream forms to stumpy forms, and from procyclic forms to bloodstream forms. The bi-stable character can be explained by the fact that RBP10 directly suppresses procyclic-specific post-transcriptional regulators. While RBP10 is present, the bloodstream-form state is self-reinforcing; loss of RBP10 enables procyclic regulators to be expressed, while expression of RBP10 in procyclic forms tips the balance towards a bloodstream-form expression pattern. In conclusion, our results demonstrate how differentiation states can be maintained by a post-transcriptional cascade.

## Materials and methods

### Parasite culture and plasmid constructs

*T. brucei* Lister 427 [67] and pleomorphic AnTat 1.1 (EATRO 1125) [56] cells expressing the Tet-repressor were used. Lister 427 were used for all experiments except those shown in Figs 5 and 6. Bloodstream form parasites were cultured in HMIi-9 medium supplemented with 10% fetal bovine serum at 37^o^C with 5% CO2. The procyclic forms were grown at 27^o^C in MEM-Pros medium supplemented with 10% heat inactivated fetal bovine serum. Stable cell lines were generated and maintained as described in [56,67] except that selection of the pleomorphic cells was done with 8µg/ml hygromycin and 2µg/ml blasticidin.

The tetracycline inducible constructs for *RBP10* RNAi and over expression were described in [37]. All new constructs and oligonucleotides are in S6 Table. A bloodstream-form cell line expressing all RBP10 with a tandem affinity purification tag at the N-terminus was generated by inserting the tag at the 5′ end of one endogenous *RBP10* open reading frame (pHD2506) and deletion of the second copy (pHD2061). Plasmids for the tethering assays are in S6 Table. The tethering constructs were separately transfected in a cell line constitutively expressing the *CAT* reporter with 5 copies of *boxB* preceding the *ACT* 3′ UTR. Expression of the fusion protein was induced for 24h using tetracycline (100ng/ml) and the CAT assays carried out as described in [25].

### Cross-linking and RNA immunoprecipitation

3x10^9^ bloodstream form cells expressing TAP-RBP10 were irradiated using UV (400 mj/cm2), washed in cold PBS and the cell pellet snap frozen in liquid nitrogen. The RNA immunoprecipitation was done as described in [24]. Briefly, the extracts were incubated with the beads, and the unbound fraction was collected. After washing, bound RBP10 was eluted using TEV protease. After digestion with 50µg proteinase K at 42°C for 15 minutes to reduce cross-linked protein, RNA was isolated from both the bound and unbound fractions using Trifast reagent (Peqlab, GMBH). To assess the quality of the purified RNA, an aliquot of the sample was analysed by Northern blotting and the blot hybridized with a splice leader probe. Total RNA from the unbound fraction was depleted of ribosomal RNA (rRNA) using RNAse H as described in [68] except that a cocktail of 50-base DNA oligos complementary to trypanosome rRNAs was used [63]. The recovered RNA from both bound and unbound samples was then analysed by RNA-Seq.

### Tandem affinity purification (TAP) and genome-wide yeast two-hybrid screen

Approximately 2x10^10^ bloodstream form cells expressing in-situ TAP-RBP10 were harvested and used for TAP as previously described [69]. Three biological replicates were done, with or without RNAse A treatment; triplicate results from a cell line with inducible GFP-TAP served as the background control. The eluate was separated on a 10% SDS-polyacrylamide gel for only 2 cm and the proteins visualized by colloidal Coomassie staining. The gel area was excised and analysed by mass spectroscopy. For the yeast two-hybrid screen, the RBP10 open reading frame was cloned into the pGBKT7 plasmid (pHD2361). The construct was used as a bait to screen a library made up of *T. brucei* random genomic DNA fragments. The screening, high throughput sequencing and analysis were done as previously described [70].

### Polysome fractionation

Approximately 3-5x10^8^ cells were collected by centrifugation (850 × g, 10 minutes, 21^o^C), and then treated in serum free media with 100µg/ml cycloheximide for 7 minutes at room temperature. The pellet was washed in 1 ml ice cold PBS, lysed in 350µl lysis buffer (20mM Tris pH7.5, 20mM KCl, 2mM MgCl2, 1mM DTT, 1000u RNasin (Promega), 10µg/ml leupeptin, 100µg/ml cycloheximide, 1× complete protease Inhibitor without EDTA (Roche), 0.2% (vol/vol) IGEPAL) by passing 15-30 times through a 21G needle, followed by centrifugation (15000 × g, 10 minutes, 4^o^C) to clear the lysate. KCl was adjusted to 120 mM and the clarified lysate loaded on top of a 4 ml continuous linear 15-50% sucrose (w/v) gradient in polysome buffer (20mM Tris pH7.5, 120mM KCl, 2mM MgCl2, 1mM DTT, 10µg/ml leupeptin, 100µg/ml cycloheximide). After 2 hours of ultracentrifugation (40000 rpm, 4^o^C; Beckman SW60 rotor), 400µl fractions were collected using Teledyne Isco Foxy Jr. gradient fractionator system and RNA was isolated using Trifast reagent (Peqlab, GMBH). In the case of RBP10 tethering, lambdaN-RBP10 was induced (24h) and the distribution of the *CAT* reporter and *alpha-tubulin* mRNAs in the collected fractions detected by Northern blotting as described in [71].

For RNASeq, bloodstream form cells plus or minus *RBP10* RNAi for 15h and, the procyclic form cells with or without RBP10-myc over-expression for 6h were used. In this case 250µl fractions were collected from each gradient and RNA was isolated after pooling the fractions into two groups; i) lighter fractions including monosomes, ii) denser fractions with at least two ribosomes. The amount of mRNA in the pooled samples was assessed by Northern blotting using splice leader RNA as probe. Also, total RNA was prepared from the input samples (~10% of total cell lysate) to quantify changes in the steady state mRNAs levels. All samples were depleted of rRNAs (as above) prior to analysis by RNA-seq.

### High throughput RNA sequencing and bioinformatic analysis

RNA-seq was done at the CellNetworks Deep Sequencing Core Facility at the University of Heidelberg. For library preparation, NEBNext Ultra RNA Library Prep Kit for Illumina (New England BioLabs Inc.) was used. The libraries were multiplexed (6 samples per lane) and sequenced with a HiSeq 2000 system, generating 50 bp single-end sequencing reads.

The quality of the raw sequencing data was checked using FastQC (http://www.bioinformatics.babraham.ac.uk/projects/fastqc), and the sequencing primers removed using Cutadapt [72]. The data was aligned to the *T. brucei* TREU 927 reference genome using Bowtie [73], then sorted and indexed using SAMtools [74]. Reads aligning to open reading frames of the TREU 927 genome were counted using custom python scripts. Analysis for differentially expressed genes was done in R using the DESeq package [75] with an adjusted p-value cut-off of 0.05.

Comparative analysis was limited to a list of unique genes modified from [9]. Gene annotations are manually updated versions of those in TritrypDB (http://tritrypdb.org/tritrypdb/), and categories were assigned manually. Statistical analysis was done in R. The 3′ UTR motif enrichment search was done using DREME [76]; annotated 3′ UTRs sequences were downloaded from tritrypDB and we considered only the mRNAs with 3′ UTR >20 nt. Manual 3′-UTR annotation was done using the RNASeq reads and poly(A) site data [9,11,77] in tritrypDB.

### Analysis of 3′-UTR reporters

Bloodstream form cells expressing the *CAT* reporter with either full length *EP1* 3′ UTR (pHD1610) or a mutant version (pHD1611) lacking the 26mer (*EP1∆26*) instability element [34] were used for RNA immunoprecipitation. For the pull down, 4x10^8^ cells (without the stem-loop) were irradiated using UV (400 mj/cm^2^), washed in cold PBS and the cell pellet lysed in 350µl lysis buffer (10mM Tris pH 7.5, 10mM NaCl, 1000u RNasin (Promega), 1× complete protease Inhibitor without EDTA (Roche), 0.1% IGEPAL) by passing 15 times through a 21G needle. The lysate was cleared by centrifugation at 15000g for 10 minutes at 4^o^C, NaCl was adjusted to 150mM followed by incubation with 50µl anti-RBP10 [37] coupled agarose beads for 2 hours at 4^o^C. After washing the beads 5 times with IPP150 buffer (10mM Tris pH 7.5, 150mM NaCl, 0.1% IGEPAL), the beads were treated with 20µg proteinase K at 42^o^C for 15 minutes and RNA was isolated from both bound and unbound fractions using Trifast reagent (peqlab, GMBH). Equal amounts of the recovered RNA (eluate and flow through fractions) were reverse-transcribed using RevertAid First Strand cDNA Synthesis Kit (Thermal Scientific) according to the manufacturer’s instructions. 2µl of the cDNA was used as template in a 50µl PCR reaction to detect the *CAT* and *alpha-tubulin* genes. PCR was done using Q5 DNA polymerase and buffer (NEB) with 0.5µm of the following primers, for *CAT* (CZ5725; CZ689) and *alpha-tubulin* (CZ5725; CZ6168); the forward primer is the same for both since it anneals to the splice leader. Aliquots (10µl) were removed after 27, 31 and 36 cycles and analysed by agarose gel electrophoresis.

To determine if the regulation of the *EP* mRNA by RBP10 depends on the 26mer instability element, a stem-loop construct (pHD1984) targeting *RBP10* was transfected into the two cell lines [37]. *RBP10* RNAi was induced using 100ng/ml tetracycline for 17h or 24h and CAT activity was measured.

### RBP10 and trypanosome differentiation

For bloodstream-procyclic form conversion, pleomorphic AnTat 1.1 bloodstream form cells with stem-loop RNAI (pHD1984) targeting *RBP10* were used. 17h after RNAi induction, the cells were pelleted, re-suspended (~8x10^5^ cells/ml) in procyclic form (MEM-pros) medium and incubated at 27^o^C. As positive controls, wild type or uninduced cells (2x10^6^ cells/ml) were treated with 6mM cis-aconitate (Sigma) at 27^o^C; after 17h the cells were transferred into procyclic form media (~8x10^5^ cells/ml) and maintained at 27^o^C. Samples were taken at different times for Western blotting (~5x10^6^) and for morphological analysis.

To convert procyclic to bloodstream forms, AniTat 1.1 bloodstream form cells with an inducible RBP10-myc construct (pHD2098) were differentiated to procyclic forms using cis-aconitate as described above. The cells were cultured in presence of hygromycin (8µg/ml) and phleomycin (0.2µg/ml) for more than 3 months to generate well-established procyclic forms. RBP10-myc was induced using 100ng/ml tetracycline. Marker proteins were detected by Western blot. VSG transcripts were detected by semi-quantitative RT-PCR as previously described [61] using primers CZ6308/CZ6309. To obtain growing bloodstream forms, RBP10-myc was induced for 48 hours, the cells were pelleted, re-suspended (2x10^5^ cells/ml) in HMI-9 medium and then incubated at 37^o^C with 5% CO_2_. The cell density was monitored for 6 days or more; wild type or uninduced cells served as control.

### Morphological analysis

To stain surface proteins, dried smears were fixed with 100% methanol at −20^o^C for 15 minutes, rehydrated in 1x PBS for 15 minutes, blocked for 20 minutes with 20% FCS, then labeled with mouse anti-EP (1:500; Cedarlane, Canada), mouse anti-phospho-GPEET (1:500; Cedarlane, Canada) and rabbit anti-VSG-117 (1:500, from either G. Cross or P. Overath). Anti-BARP staining was done as described in [5], using a procyclic cell line with the *BARP* open reading frame in the procyclin locus as a positive control. For PGK staining, 2x10^6^ cells were collected by centrifugation, re-suspended, fixed in 4% paraformadehyde (in 1x PBS) for 18 minutes, sedimented again for 2 minutes, re-suspended in PBS and allowed to settle on poly-L-lysine coated slides for 30 minutes. Before staining, slides were blocked with 20% fetal calf serum (1x PBS) for 20 minutes. Cells were permeabilised with 0.2% (v/v) Triton-X 100 (in 1xPBS) for 15 minutes at room temperature, washed twice then incubated for one hour with rabbit anti PGK antibody (1:1500, [78]). The cells were washed 3 times before being stained with fluor-conjugated secondary antibody (1:500; mouse-Cy3 or rabbit-Alexa-488; Molecular probes, Eugene). Cellular DNA was stained with 100ng/ml DAPI in 1x PBS for 15 minutes. Images were taken using Olympus Cell-R microscope. Random fields were photographed (by E.M.) and analysed (by C.C.) using Fiji [79]. We attempted blind analysis but this was not possible because the differences between samples were too obvious.

### Flow cytometry

Approximately 5x10^6^ cells were fixed with 2% formaldehyde/0.05% glutaraldehyde at 4^o^C for at least 1 hour. The cells were pelleted, washed twice with PBS then incubated with 200µl (2% BSA in PBS) mouse anti-EP (Cedarlane, Canada; 1:500), anti phosph-GPEET (Cedarlane, Canada; 1:500), or rabbit anti-VSG-117 (1:500) for one hour on ice. After washing twice, the cells were stained with the secondary antibody (1:500; mouse-Cy5 or rabbit-Alexa-488; Molecular probes, Eugene) for one hour. Cells stained only with the secondary antibody and the unstained cells served as the negative controls. Flow cytometry was performed with BD FACSCanto II flow cytometer and the data analysed using FlowJo software (TreeStar Inc.).

### Western blotting

Western blotting was done as previously described [71]. Antibodies were: anti-RBP10 (rat, 1:2000, [37]), anti-HNRNPH (rabbit,1:5000, [20]), anti-TAO (rabbit,1:100, [78]), anti-EP (mouse,1:2000, Cedarlane, Canada), anti-phospho-GPEET (mouse,1:2000, Cedarlane, Canada), anti-BARP (rabbit,1:2500, [5]), and anti-PIP39 (rabbit,1:1000, [55]) and anti-PAD1 (rabbit,1:1000, [54] with electrophoresis under non-denaturing conditions).

### Data availability

All sequence datasets are available at Array Express with the accession number E-MTAB-4564.

### Ethical approval

The research did not involve animals or humans and did not require ethical approval.

## Acknowledgements

For antibodies we thank Keith Matthews (Edinburgh University, Scotland; PIP39 and PAD1), Isabel Roditi (University of Bern, Switzerland, BARP); Shula Michaeli (Bar-Ilan University, Israel, anti-HNRNPH), Paul MIchels (anti-PGK, Edinburgh University, Scotland) and Minu Chauduri (Meharry Medical College, Nashville, Tennessee, USA, AOX). The anti-VSG117 antibody was either from Peter Overath (Max-Planck Institut, Tübingen, Germany) or George Cross (Rockefeller University, USA). We also thank Keith, Isabel and Shula for useful discussions. All RNA-Seq libraries were prepared by David Ibberson at Bioquant, Heidelberg, and sequenced at EMBL. We thank Claudia Helbig and Ute Leibfried for technical assistance, Dorothea Droll for the 4EIP yeast 2-hybrid constructs, and Kevin Leiss for counting the numbers of elements in all 3′-UTRs.

## Supporting information

### S1 Text

3′-untranslated regions of mRNAs that were at least 3x enriched in both RBP10 pull-downs.

### S2 Text

3′-untranslated regions of mRNAs that were less than 0.7x enriched in both RBP10 pull-downs.

### S3 Text

3′-untranslated regions of mRNAs that were at least 3x enriched on average in RBP10 pull-downs, according to DESeq (Padj <8 E-5), but one replicate was less than 3x enriched.

### S1 Table

Proteins that co-purified with TAP-tagged RBP10.

Endogenously TAP-tagged RBP10 was purified three times in the presence or absence of RNase. The controls were three independent purifications of GFP-TAP

Sheet 1: Proteins enriched in the purifications. Peptide counts and ratios are shown. The highest peptide count from the GFP-TAP replicates was used as a baseline. The proteins shown in this table were present as at least 2 peptides, present in at least two similar purifications, and were on average at least 2-fold enriched relative to GFP-TAP. Functions of interest are highlighted in different colours. “Average PGKB =tet/-tet” are tethering screen results [39]; a value above 1.5 suggests that the protein can suppress expression. “in starvation granules?” is 1 if positive and 0 if negative [80]. The number of peptides found by purification of MKT1-TAP are also indicated [70].

Sheet 2: All raw data.

### S2 Table

Results of a yeast 20-hybrid screen with a random fragment library.

The library [70] was screened with pGBKT7-RBP10 and results evaluated by sequencing.

Sheet 1: Raw data for pGBKT7-RBP10. All reads that started from position −36 onwards (relative to the ATG) are included.

Sheet 2: Raw data for pGBKT7 (unselected background, filtered for at least 4 reads)

Sheet 3: Results for pGBKT7-RBP10, filtered to exclude loci with less than 10 reads, and open reading frames with less than 2 independent fragments. Numerous cysteine protease fragments have been removed.

Sheet 4: Possible interaction partners. The ratio to the maximum background count is at least 4. Proteins that also scored as interacting with MKT1 or CFB1 [70] (indicating less specificity) are listed separately. The “regulator” column shows results from the tethering screen [39].

### S3 Table

Effect of RBP10 depletion on gene expression in bloodstream forms.

RNAi was induced for 15h and mRNA was collected for RNASeq from total cells, the free and monosomal fractions, and the polysomal fractions. Data are for two independent experiments which are labelled A and B.

Throughout the Tables, the colour code for ratios is as follows: Red is >2, orange is 1.5-2.0, green is 0.667 - 0.5 and blue is <0.5. Shading of columns does not follow a particular colour code but is designed to make the tables easier to navigate. Many numbers have been rounded to 2 or 3 points beyond the decimal in order to decrease the sizes of the files. Ribosome profiling results (Riboprof) are from [11], and developmental regulation data for total RNA are from [10].

Sheet 1: Raw read counts for all unique genes, chosen according to [9] with some later additions.

Sheet 2: Columns E - P Results were converted to reads per million reads (RPM). For each experiment, we then corrected the RPM for the monosomes and polysomes according to the fraction of the total mRNA that was in each fraction (sheet 7 and S3 Fig, numbers not included in the Table). We then calculated the percentage of each mRNA that was on polysomes (columns Q-T). Column U contains the average of ratio between RNAi and control, calculated for the total input RNA - (K+L) / (E+F). Column V is the same ratio for the fraction of RNA not on polysomes, and column W is the ratio for the polysomal fractions (S+T) / (Q+R). To find the lowest possible estimate of the ratio in W, we took the lowest value for the proportion in polysomes after *rbp10* RNAi, and divided by the highest value from wild-type. From this column (X) we could find mRNAs that reliably move *into* polysomes after RNAi. Column Y is similar but shows the *highest* estimate for the ratio; this can be used to find mRNAs that reliably move *away* from polysomes after RNAi. Since there were only 3 genes in this category (column Y) they were not considered further.

Sheet 3: mRNAs that move onto polysomes after *rbp10* RNAi. Same as sheet 2, except that only genes for which column W is more than 1.5, and column X is more than 1.2, are included. (This means that the average increase is at least 1.5x and the minimum is 1.2x.) Columns Y and Z are ribosome profiling results from [11], giving the ribosome density ratios for bloodstream forms (BS) divided by procyclic forms (PC). Slender bloodstream forms from blood are slBS and cultured bloodstream forms are cBS. Column AB is results from S5 Table, sheet 2 (binding of the RNA to RBP10, ratio of bound to unbound) and column AC is the adjusted P-values for the results in column AB.

Sheet 4: DeSeq analysis of the data in Sheet 1.

Sheet 5: Genes showing at least a 1.5-fold increase either in the amount of total RNA, or the amount of RNA in polysomes, or both, according to DeSeq (adjusted P-value threshold 0.05). Column AA is the RNA-binding ratio from S5 Table, sheet 2,

Sheet 6: As sheet 5, except that these are genes showing at least a 1.5-fold decrease either in the amount of total RNA, or the amount of RNA in polysomes, or both, according to DeSeq (adjusted P-value threshold 0.05).

Sheet 7: Quantitation of spliced leader signals for the free/monosome and polysome fractions used for RNA-Seq

Sheet 8: Statistical analyses to compare the proteomes of differentiating trypanosomes [16] with the effects of RBP10 RNAi on RNA levels.

### S4 Table

Effect of RBP10 expression on gene expression in procyclic forms.

Expression of RBP10 was induced with tetracycline addition for 6 hours, and RNA samples prepared from duplicates as in Supplementary Table S3. Controls are the same cells without tetracycline.

Sheet 1: Columns D-O are raw reads. Columns P-S are corresponding RPM values for input samples.

Sheet 2: Proportions in free/monosome and polysome fractions were calculated as described for Supplementary Table S3, sheet 2. Columns Q-X: Effects on both polysomes and free/monosome fractions were also calculated in a similar way.

Sheet 3: As sheet 2, but only genes showing regulated translation according to the following criteria: (a) movement to the free/monosome fraction from the polysomes. This is the top section (rows 2-320). The smallest value for the proportion in free/monosomes with RBP10 expression was divided by the largest value calculated value for the proportion in free/monosomes without RBP10 induction. If this number was greater than 1.5, the gene was included. In addition, we divided the largest number for the proportion in polysomes plus RBP10 expression by the smallest value for the proportion in polysomes without RBP10 induction. If this number was less than 0.667 the gene was also included. Genes that satisfied both criteria are in bold. Additional columns are as in Table S3. Rows 322-326 are the four RNAs showing movement onto polysomes.

Sheet 4: DeSeq analysis of the results in Sheet 1.

Sheet 5: Genes showing at least a 1.5-fold increase either in the amount of total RNA, or the amount of RNA in polysomes, or both, according to DeSeq (adjusted P-value threshold 0.05).

Sheet 6: As sheet 5, except that these are genes showing at least a 1.5-fold decrease either in the amount of total RNA, or the amount of RNA in polysomes, or both, according to DeSeq (adjusted P-value threshold 0.05). Columns V-Z are results for salivary gland trypanosomes, calculated from the raw data in [38]; statistical analysis was not possible due to the absence of replicas.

Sheet 7: Quantitation of spliced leader signals for the free/monosome and polysome fractions used for RNASeq.

### S5 Table

RNAs bound to tagged RBP10.

Results for two independent experiments, A and B, are shown.

Sheet 1: Raw read counts for all samples, calculation of reads per million reads (RPM), and ratios of bound/unbound for each experiment.

Sheet 2: DeSeq analysis of the raw data in sheet 1

Sheet 3: Set of mRNAs that were at least 3-fold bound according to DeSeq. This is a slightly more relaxed stringency than that used in sheet 5. The largest P-value was 7E-5. H-J: Results for developmental regulation [10]; K - calculated from [38]; L-T: results from Supplementary Tables S3 and S4; U, W, Z: [13]; V: [14]; X, Y, AA: [15]; AB: [16]; AC: [81]; AD: [42].

Sheet 4: Statistics used to create the Venn diagrams in Figure 2.

Sheet 5: Set of bound RNAs used for the motif search. Filters were (a) at least 10 reads in all samples and (b) a bound/unbound ratio of >3 in both experiments. The 3′-UTR lengths are those annotated in TritrypDB, except where noted. Only annotated 3′-UTRs were used for the initial motif search.

### S6 Table

Plasmids and oligonucleotides

**S1 Fig.** Alignment of RBP10 from different kinetoplastids

**S2 Fig.**
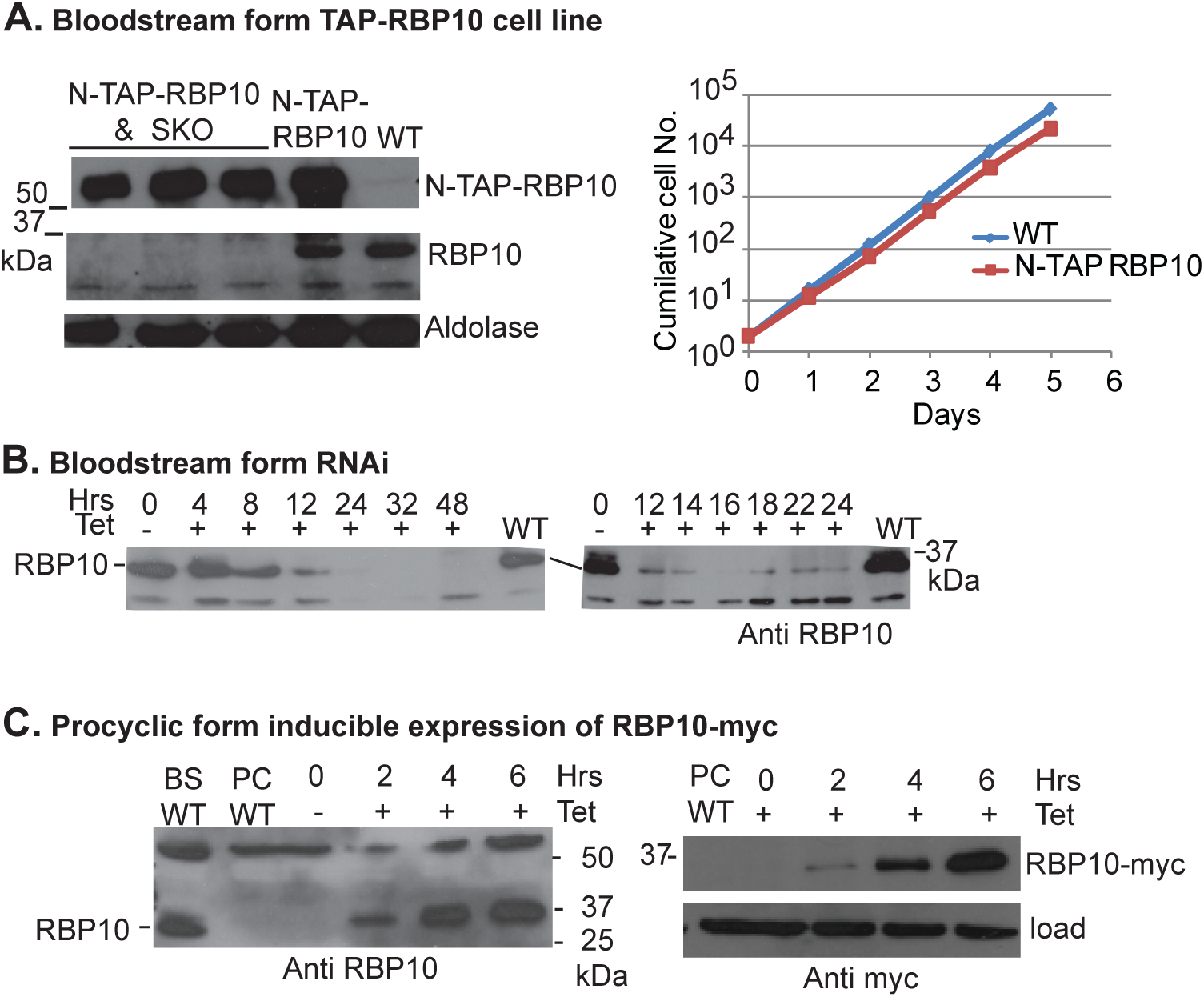
Cell line characterisation and RBP10 expression A) Characterisation of bloodstream-form trypanosomes with a sequence encoding an N-terminal tandem affinity purification tag integrated in frame with the *RBP10* coding region. The left hand panel is a western blot showing the presence or absence of TAP-RBP10 and native RBP10, with genotypes above the lanes. The right-hand panel is a cumulative growth plot for the cell line used in the interactome experiments. B) RNAi cell line: Western blots showing the time course of RBP10 decrease after tetracycline addition C) Western blots showing the time course of expression of RBP10-myc in Lister 427 procyclic forms after tetracycline addition.

**S3 Fig.**
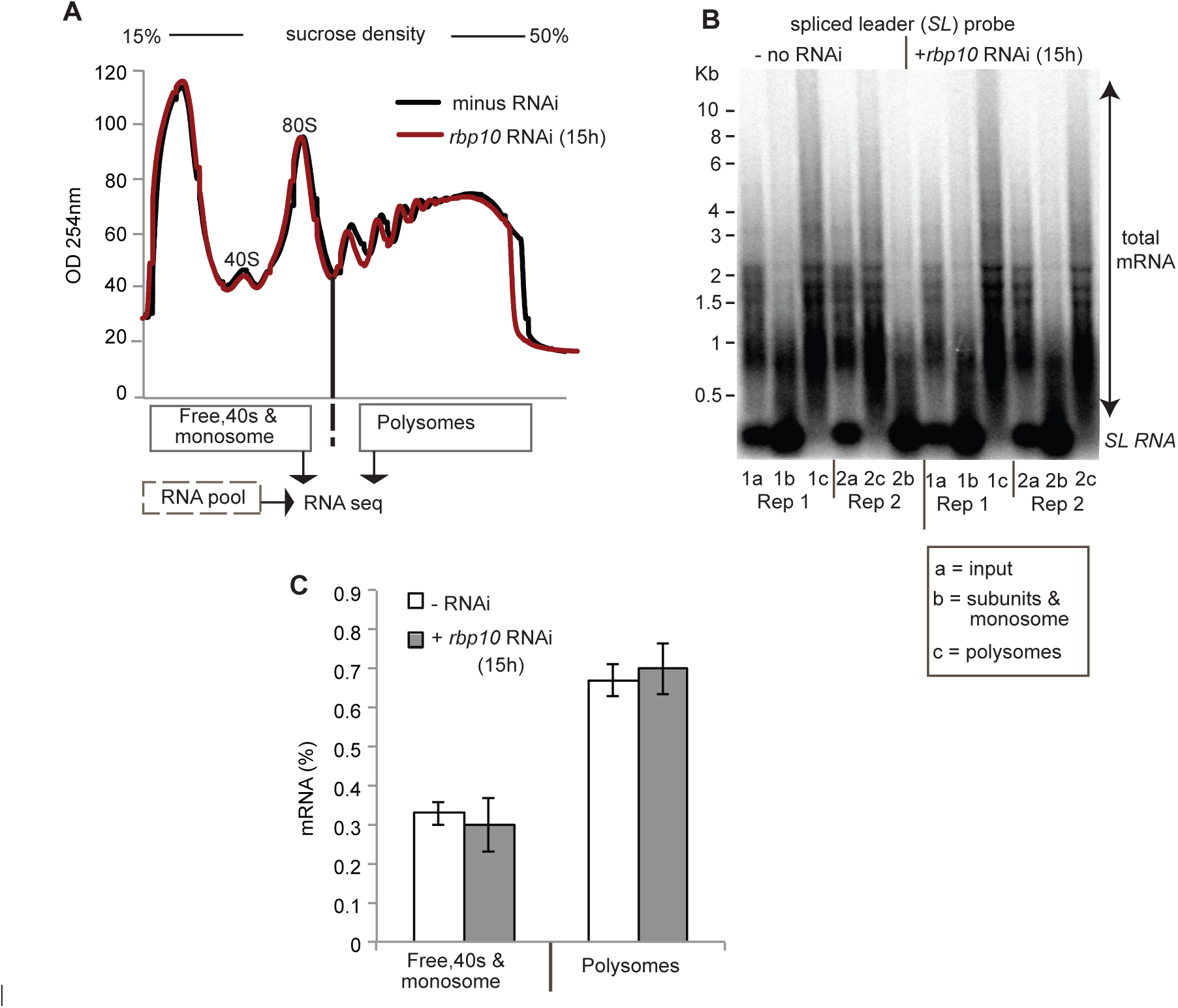
RNAi targeting *RBP10* in bloodstream forms: samples used for RNASeq A. Typical Sucrose gradient profiles of samples B. Total mRNA in the sucrose gradient fractions, detected using the spliced leader probe C. Average quantitation of the spliced leader signal

**S4 Fig.**
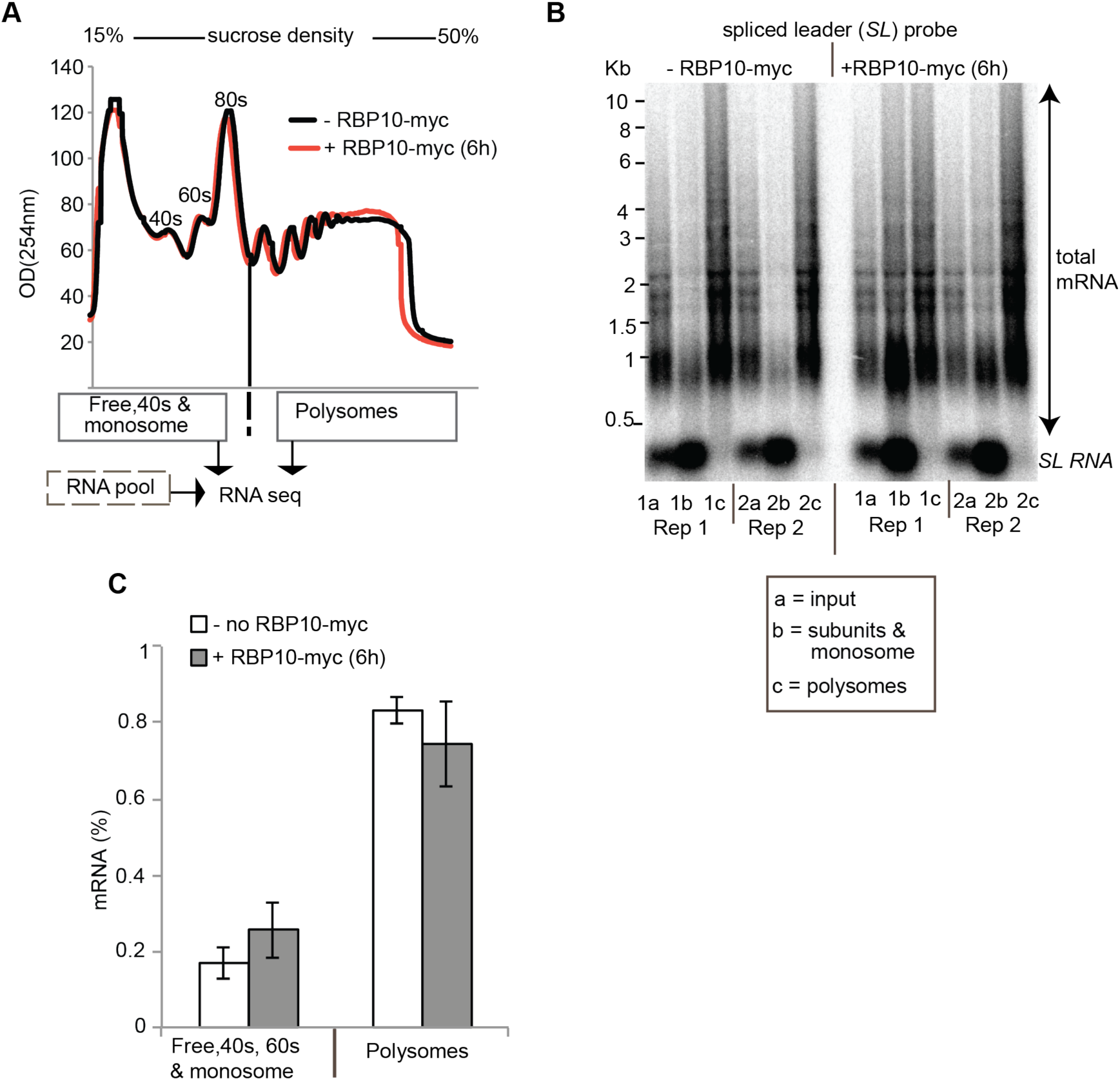
Expression of RBP10 in procyclic forms: samples used for RNASeq A. Typical Sucrose gradient profiles of samples B. Total mRNA in the sucrose gradient fractions, detected using the spliced leader probe, C. Average quantitation of the spliced leader signal

**S5 Fig.**
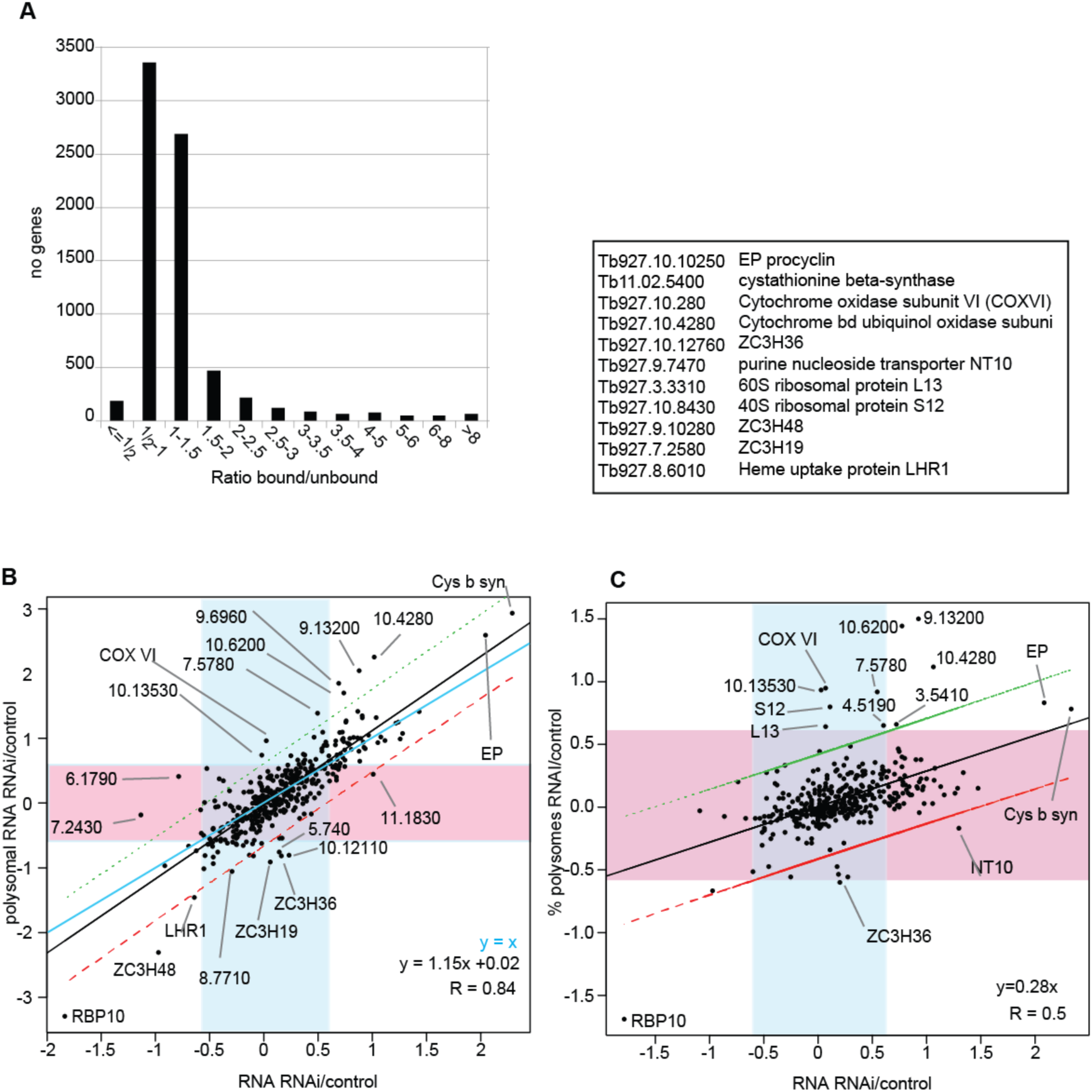
Effects of RNAi and analysis of RBP10-bound RNAs. A. Analysis of binding of individual mRNAs to RBP10. The classes are ratios of bound/unbound; the number of different open reading frames in each class is on the y axis. B. Scatter plot comparing the effect of *rbp10* RNAi on total RNA (x axis) with the effect on polysomal RNA (y axis) for mRNAs that were at least 3x enriched according to DeSeq; all P-values were less than 8E-5. The black line is the regression line and the red and green lines show the 95% confidence limits for the data. The blue shadow encloses total mRNAs that were less than 1.5x affected, and the pink shadow encloses polysomal RNAs that were less than 1.5x affected. The cyan line is perfect correlation. The box beneath the graph lists relevant TritrypDB accession numbers. The gene numbers on the plot are also accession numbers, with “Tb927.” removed. C. As (B), but here the y axis shows the effect of RNAi on the percentage of the mRNA in polysomes.

**S6 Fig.**
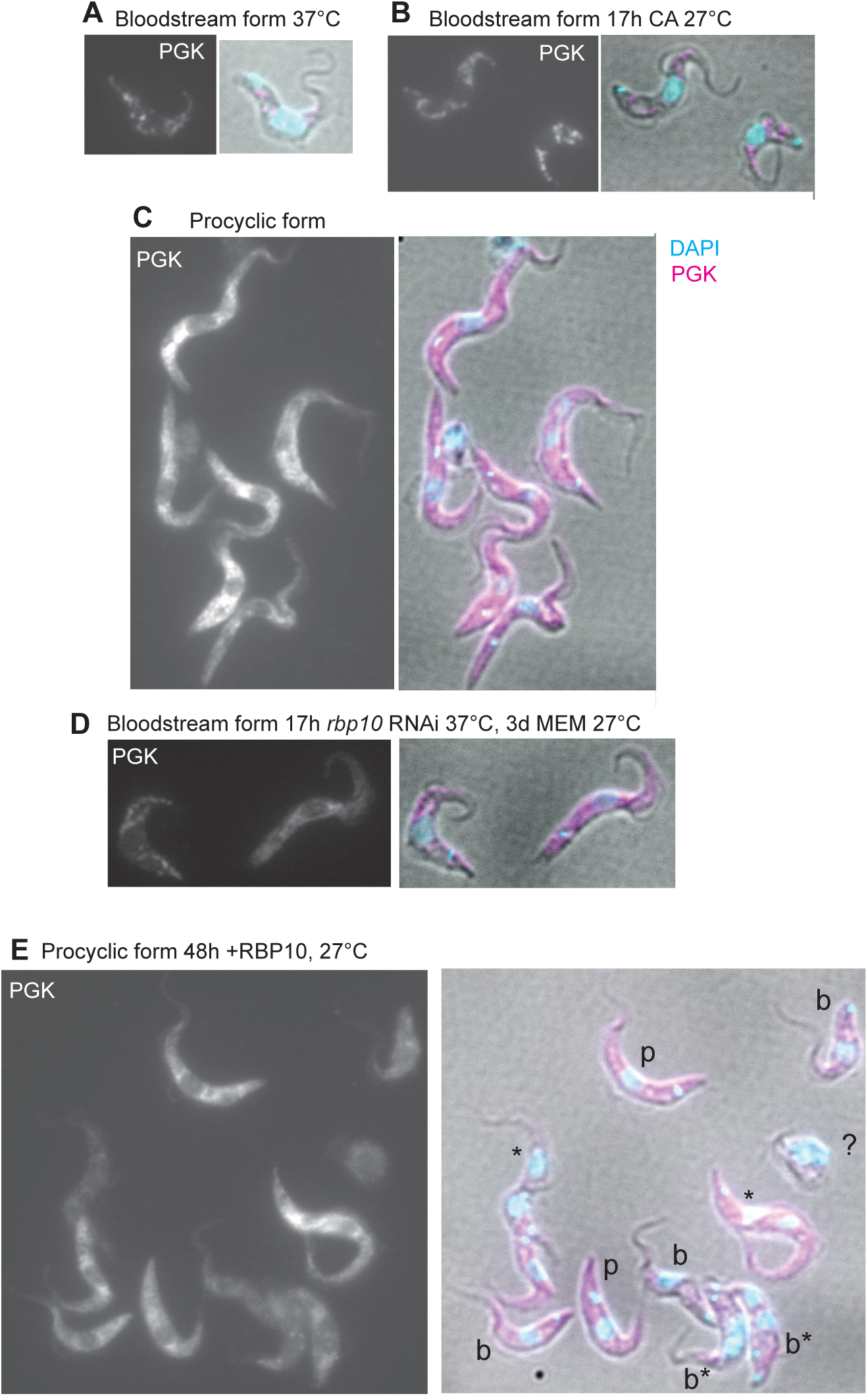
Relocation of phosphoglycerate kinase during differentiation. Cells were fixed with fomaldehyde, pernmeabilized with triton x-100, and stained using a polyclonal antibody to phosphglycerate kinase (PGK). The grey-scale panels show PGK alone and the differential interference contrast panels show DNA in cyan and PGK in magenta. A. Bloodstream forms B. Bloodstream forms after incubation with cis aconitate for 17h at 27°C. C. Procyclic forms. D. Bloodstream forms with 17h *rbp10* RNAi followed by culture for 3 days under procyclic-form culture conditions. The selection shows on procyclic-like trypanosome (left) and one which still has bloodstream-form morphology (right). E. Procyclic forms after 2 days induction of expression of RBP10-myc. Cells with terminal kinetoplasts are labelled “b” and cells with more procyclic morphology are labelled “p”. Cells that appear to be dividing are indicated with asterisks. A very abnormal cell is indicated with “?”. Cells if unclear status are not labelled.

**S7 Fig.**
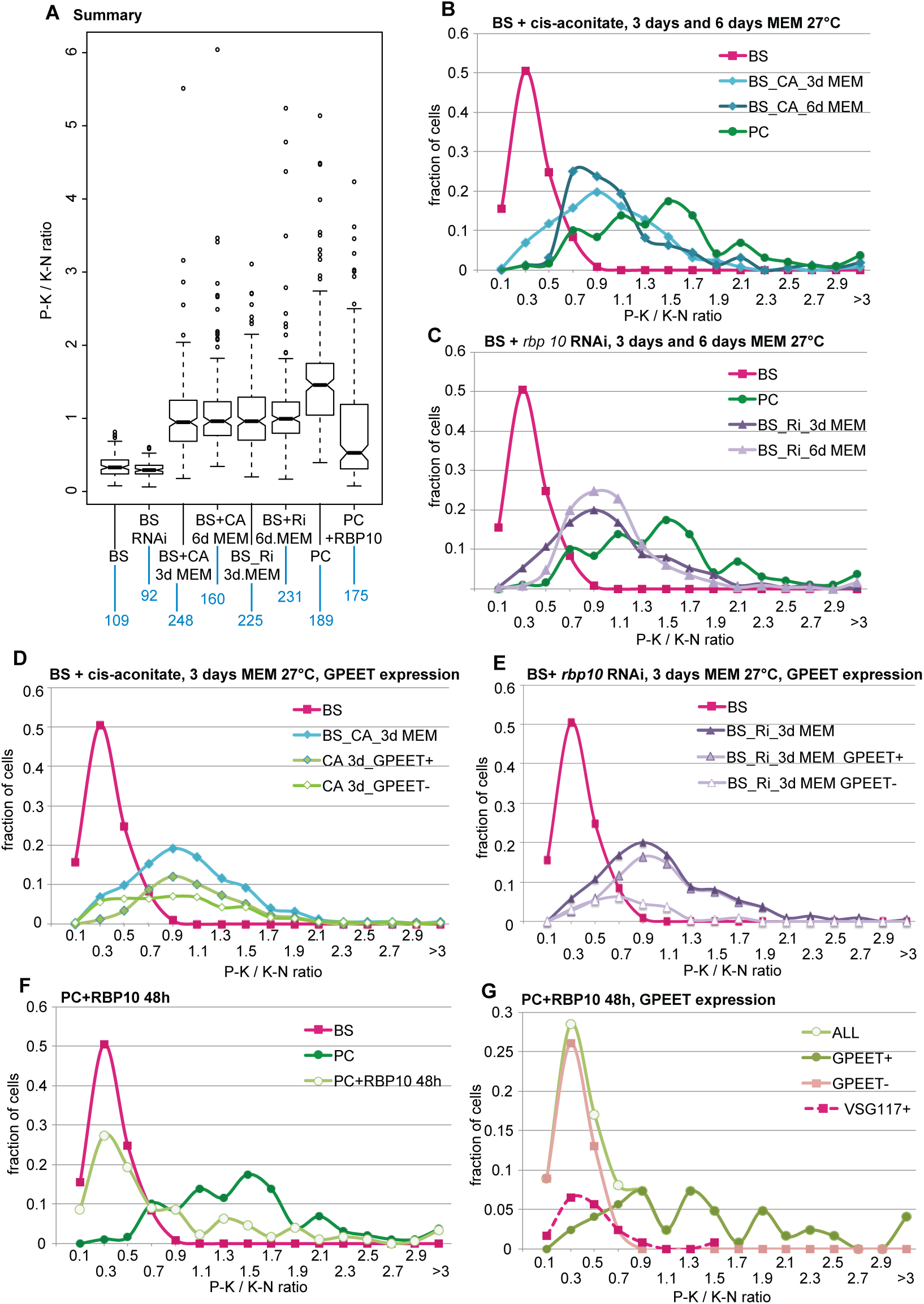
Morphological analysis of differentiation. Cells were treated as indicated, fixed, and stained for DNA (cyan), EP-procyclin, phopho-GPEET procyclin, or VSG. (magenta). The posterior-kinetoplast and kinetoplast-nucleus distances were measured in Image J and the ratios calculated. A. Summary of all results. This is the same as Fig 5H, but all of the outliers are included, and the numbers of cells analysed are below each lane. BS - normal bloodstream forms; PC-normal procyclic forms; CA - 17h cis-aconitate pretreatment; RNAi or Ri; 17h *rbp10* induction; +RBP10 - induced expression or RBP10 for 2 days. B. Cell distributions after cis-aconitate-induced differentiation. The x axis shows the P-K/K-N ration, and y-axis shows the percentage of cells with that ratio. Controls are bloodstream forms (magenta) and established procyclic forms (green). The other lines are for 3 and 6 days after transfer to procyclic medium (MEM). C. As B, but with transfer after 17h *rbp10* RNAi. D. Phospho-GPEET expression 3 days after cis-aconitate-stimulated differentiation. The cells indicated with the cyan line (BS_CA_3d MEM) were subdivided into GPEET positive and GPEET negative. E. Phospho-GPEET expression 3 days after RNAi-stimulated differentiation; other details as in D. F. Cell distributions in procyclic forms after induced expression of RBP10. G. As (F), but also showing staining with anti-phospho-GPEET and anti-VSG117. Cells wit no GPEET have bloodstream-form kinetoplast positions, and a subset of them cross-reacts with anti-VSG117 antibodies. The remainder may express VSGs that do not cross react with the anti-117 antibody.

